# Associations between music and dance relationships, rhythmic proficiency, and spatiotemporal movement modulation ability in adults with and without mild cognitive impairment

**DOI:** 10.1101/2023.12.19.572238

**Authors:** Alexandra Slusarenko, Michael C. Rosenberg, Meghan E. Kazanski, J. Lucas McKay, Laura Emmery, Trisha M. Kesar, Madeleine E. Hackney

## Abstract

**Background:** Personalized dance-based movement therapies may improve cognitive and motor function in individuals with mild cognitive impairment (MCI), a precursor to Alzheimer’s disease. While age- and MCI-related deficits reduce individuals’ abilities to perform dance-like rhythmic movement sequences (RMS)—spatial and temporal modifications to movement—it remains unclear how individuals’ relationships to dance and music affect their ability to perform RMS.

**Objective:** Characterize associations between RMS performance and music or dance relationships, as well as the ability to perceive rhythm and meter (rhythmic proficiency) in adults with and without MCI.

**Methods:** We used wearable inertial sensors to evaluate the ability of 12 young adults (YA; age=23.9±4.2 yrs; 9F), 26 older adults without MCI (OA; age=68.1±8.5 yrs; 16F), and 18 adults with MCI (MCI; age=70.8±6.2 yrs; 10F) to accurately perform spatial, temporal, and spatiotemporal RMS. To quantify self-reported music and dance relationships and rhythmic proficiency, we developed Music (MRQ) and Dance Relationship Questionnaires (DRQ), and a rhythm assessment (RA), respectively. We correlated MRQ, DRQ, and RA scores against RMS performance for each group separately.

**Results:** The OA and YA groups exhibited better MRQ and RA scores than the MCI group (p<0.006). Better MRQ and RA scores were associated with better temporal RMS performance for only the YA and OA groups (r^2^=0.18-0.41; p<0.045). DRQ scores were not associated with RMS performance in any group.

**Conclusions:** Cognitive deficits in adults with MCI likely limit the extent to which music relationships or rhythmic proficiency improve the ability to perform temporal aspects of movements performed during dance-based therapies.

## Introduction

Dance-based movement therapy is a cognitively engaging physical activity that helps mitigate neurodegeneration and improve cognitive and motor function in individuals with mild cognitive impairment (MCI), a precursor to Alzheimer’s disease and dementia [1-5]. The level of motor and cognitive challenge of dance-based therapies may be customized by selecting different therapy parameters (e.g., prescribed dance movements or musical elements). Selecting therapy parameters that challenge each individual, without being discouragingly difficult, may enhance therapeutic efficacy [3, 6]. However, we currently lack objective approaches to personalize dance-based therapy parameters [1, 7]. The ability to perform dance-based therapies likely depends on multiple factors, including aspects of motor and cognitive function, as well as each individual’s relationships to music and dance (defined as individuals’ histories, experiences, and attitudes towards music and dance) and rhythmic proficiency (defined as individuals’ abilities to perceive and replicate rhythms) [1, 3, 5, 7]. While we previously showed that age-related declines in motor and cognitive function reduce the ability to perform dance-like movements, it remains unclear how relationships with dance and music, or rhythmic proficiency, impact this ability [8].

Assaying an individual’s ability to perform dance-like movements is critical for determining challenging, engaging, and individual-specific dance therapy protocols. We recently developed a library of Rhythmic Movement Sequences (RMS) that isolate spatial (e.g., modified lower-extremity joint range of motion and coordination) and temporal (e.g., modified timing of stepping patterns, synchronized to music) features of forward movement (*i.e.*, walking) [8, 9]. Note that “RMS” used throughout the manuscript refers to the movement sequences, not “root-mean-square” error. Similar to RMS, dance therapies often involve modifications to spatial and temporal aspects of movement, but lack principles to determine modification difficulty based on participants’ abilities [3, 5, 10]. By requiring participants to walk while modifying either spatial or temporal aspects of movement, or both simultaneously, the RMS experimental paradigm provides insights regarding participant’s abilities to modify spatial and temporal aspects of movement that may be performed during therapy.

RMS performance can be quantified as the ability to achieve prescribed spatial and temporal targets [8]. Spatial RMS consist of modifications to the joint kinematics of typical walking to achieve prescribed spatial targets (e.g., 90 degrees of peak hip flexion during the swing phase of gait), with no prescribed changes in step timing [11]. Deviations from spatial modification targets may reflect an inability to perceive, comprehend, or recall spatial patterns and execute appropriate motor commands to accurately modulate spatial aspects of movement. Temporal RMS involve modifying step timing to achieve prescribed patterns of quick and slow steps, synchronized to perceived concurrent rhythmic cues in music, with no prescribed spatial modifications [9]. Our approach to temporal RMS is grounded in principles of music theory that suggest that auditory cues can influence the temporal progression of movement [12-14]. Deviations from the prescribed tempo or step pattern reflect an inability to perceive musical cues and recall or execute stepping patterns. Spatiotemporal RMS assays the additional challenge of simultaneously performing spatial and temporal RMS. The performance of these three types of RMS may be differentially influenced by multiple factors including motor function, cognitive function, and experience perceiving rhythms and executing motor commands.

We previously showed detrimental effects of age and MCI on individuals’ abilities to perform RMS, though these factors did not comprehensively explain individual differences in RMS performance [8]. We found worse performance on spatial, but not temporal RMS in older adults without MCI, compared to young adults, suggesting that age-related motor deficits are primarily related to a reduced ability to accurately modulate spatial features of movement. Conversely, we found worse performance on some spatial and some temporal RMS in older adults with MCI, compared to older adults without MCI, suggesting that cognitive deficits in adults with MCI are related to a reduced ability to accurately modify both spatial and temporal features of movement. However, we found that RMS performance was not fully explained by differences in age or cognitive status [8]. Individual differences in the ability to perceive meter and rhythm and synchronize motor commands to music may help explain variability in the temporal aspects of RMS performance in individuals of similar age and cognitive status [14, 15].

Similarly, individual differences in motor experiences stemming from music or dance relationships may explain variability in the spatial aspects of RMS performance with similar age-related motor function levels [11, 15].

Relationships to music and dance reflect individuals’ diverse histories, perceptions, and attitudes towards music and dance, while rhythmic proficiency reflects the ability to entrain motor commands to perceived rhythms. Merely by existing within a cultural context and through past experiences, individuals develop such relationships with these ubiquitous and often interconnected art forms [14, 16]. Lifelong exposure to music and dance may alter the neural and biomechanical processes underlying the ability to perceive auditory cues and produce corresponding rhythmic motor commands, both of which are required during dance-based therapies [3, 11, 15, 17, 18]. Stronger relationships with music may enhance the ability to perceive and predict the rhythmic patterns, musical groupings, and meter needed to follow temporal movement cues in music [15, 19, 20]. Stronger relationships with dance may enhance the ability to sense and accurately modify spatial aspects of movement (e.g., joint kinematics) and entrain movements to musical cues [11, 15]. Conversely, stronger rhythmic proficiency reflects a better ability to perceive rhythm and meter, anticipate beats in music, and entrain motor commands to prescribed rhythmic cues [14]. Rhythmic proficiency, therefore, reflects both perceptual acuity and motor skill pertinent to RMS performance, making it distinct from music and dance relationships [21].

Here, we investigated how individuals’ relationships with dance and music, and their rhythmic proficiency impacted their performance on spatial, temporal, and spatiotemporal RMS. The central hypothesis of this work is that stronger relationships and past experiences with music and dance, and rhythmic proficiency, contribute to an improved ability to accurately modulate spatial and temporal aspects of movement, as captured during RMS. To test this hypothesis, we developed Music Relationship (MRQ) and Dance Relationship Questionnaires (DRQ) that survey individuals’ relationships to music and dance, respectively. We also developed a Rhythm Assessment (RA) that assesses rhythmic proficiency, defined by the ability to recognize meter in music and reproduce auditory or visual rhythms. First, we characterized the relationships between the MRQ, DRQ, and RA. We then compared these novel assessment scores to RMS performance in younger adults (YA group) and older adults without (OA group) and with MCI (MCI group). We predicted that, within each group, stronger relationships to music (higher MRQ scores) would be associated with more accurate performance of temporal and spatiotemporal RMS. We similarly predicted that, within groups, stronger relationships to dance (higher DRQ scores) would be associated with more accurate performance of spatial and spatiotemporal RMS. Finally, we predicted that better rhythmic proficiency (higher RA scores) would be associated with more accurate performance of temporal and spatiotemporal RMS. Furthermore, to better understand how music-related perceptual acuity and motor skill impact RMS performance, we conducted an exploratory correlational analysis regressing two RA subcomponents—*auditory rhythm reproduction* and *auditory meter recognition*—against RMS performance.

## Materials and Methods

This study was approved by the Emory University Institutional Review Board (STUDY00003507). All participants provided written, informed consent before participation.

### Participants

An observational cross-sectional study with 56 participants was performed in non-disabled younger adults (YA; N=12), older adults without MCI (OA; N = 26), and older adults with MCI (N = 18; Table 1). The inclusion criteria for all participants were the ability to walk 20m without an assistive device, 6 years of education (e.g., kindergarten through fifth grade), proficiency in the English language, and no hospitalizations in the last 60 days. YA participants included in the study were 18-35 years of age and OA and MCI participants were 55 years and older. For participants with MCI, additional inclusion criteria included amnestic MCI, as defined using the Alzheimer’s Disease Neuroimaging Initiative criteria and standard clinical assessments showing reduced executive function, working memory, and spatial cognition [22]. Assessments characterizing cognitive function included the Montreal Cognitive Assessment (MoCA), Reverse Corsi Blocks, Body Position Spatial Task, and Trail Making Test (Table 1) [23-26]. Table 1 shows the average (±1 SD) participant demographic information and clinical assessment scores for each group, along with p-values reflecting the probability of statistical differences across the groups (omnibus ANOVA or Chi-Squared tests, where appropriate; α = 0.05). When we identified significant differences across groups, we performed post-hoc independent-samples t-tests to identify differences between groups the YA and OA groups (daggers in Table 1; α = 0.05) and the OA and MCI groups (double-daggers in Table 1; α = 0.05).

**Table 1:**
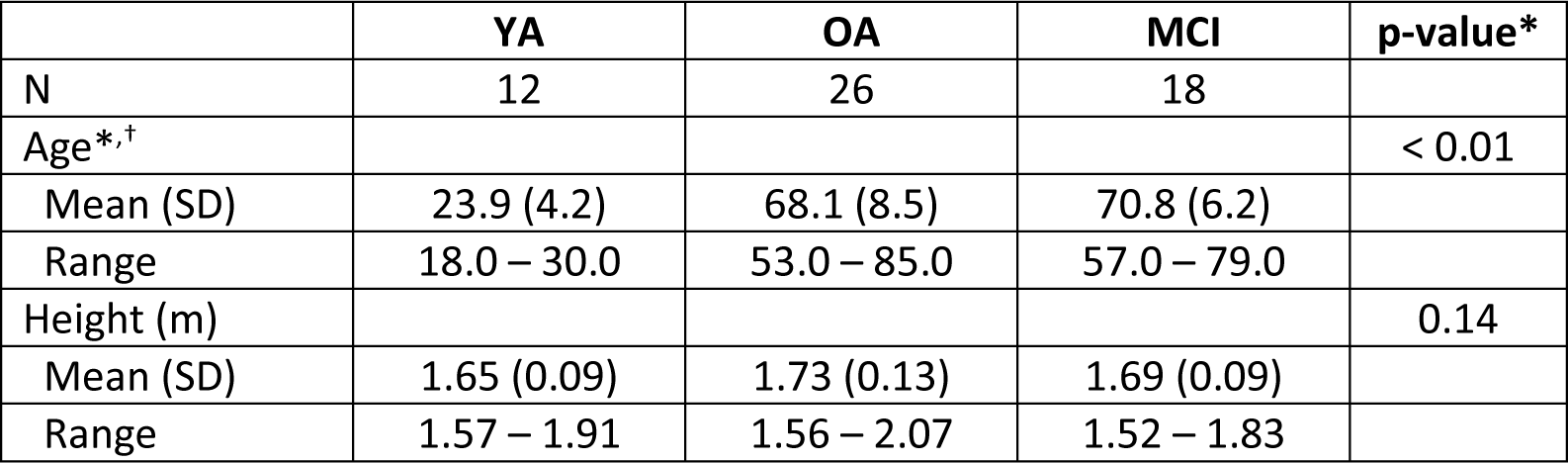
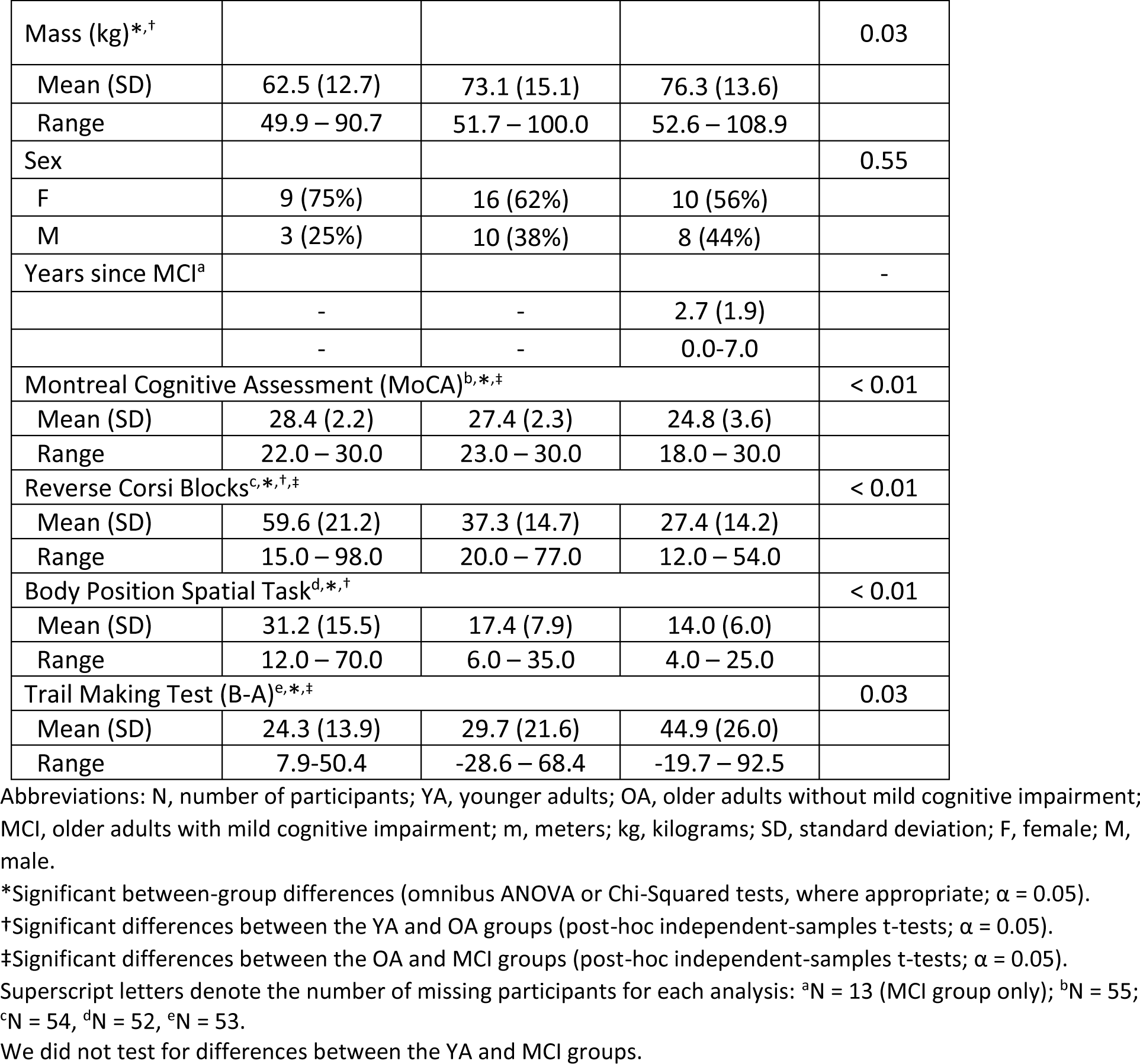
Demographic characteristics for each participant group.

### Rhythmic Movement Sequences (RMS)

Participants performed an overground assessment battery of 9 *spatial*, 9 *temporal*, and 4 *spatiotemporal* RMS, using the same protocol as described by Rosenberg and colleagues (2023) [8]. Each RMS consisted of spatial and/or temporal modifications to movement: Spatial RMS involved modifying leg joint angles during the stance or swing phases during forward movement (i.e., walking), or during both phases of gait. While dance does not always prescribe strict joint angles, and can focus instead on movement fluidity or interpersonal interactions, control of one’s kinematic configuration remains an essential aspect of some dance styles and therapies [5, 7, 17, 27]. Each spatial RMS consisted of two different spatial *gait modifications* to leg joint kinematics, with one modification per leg or both modifications in the same leg. Spatial modifications did not prescribe step timing or rhythms. The spatial modifications used in this study have corollaries in ballet, alter the typical leg joint flexion-extension patterns of walking, and were designed to be feasible for OA and individuals with MCI [28]. We defined three sub-classes of spatial RMS: *swing*, *stance*, and *swing-stance* (Figure 1). RMS sub-class names were defined by whether participants were instructed to modify joint kinematics during swing, stance, or both phases of gait, respectively. Each sub-class contained three different RMS. For *swing* and *stance* RMS, different gait modifications were performed by each leg. For example, the *Attitude-Developpé* RMS involved performing the *Attitude* and *Developpé* modifications during the swing phases of the left and right legs, respectively (Figure 1; bullets denote target joint angles). Similarly, for *swing-stance* RMS, two different modifications were performed by the left leg, while the right leg could move freely. For example, the *Attitude-Relevé* modification involved performing the *Attitude* and *Relevé* modifications both with the left leg during the swing and stance phases of gait, respectively.

**Figure 1:**
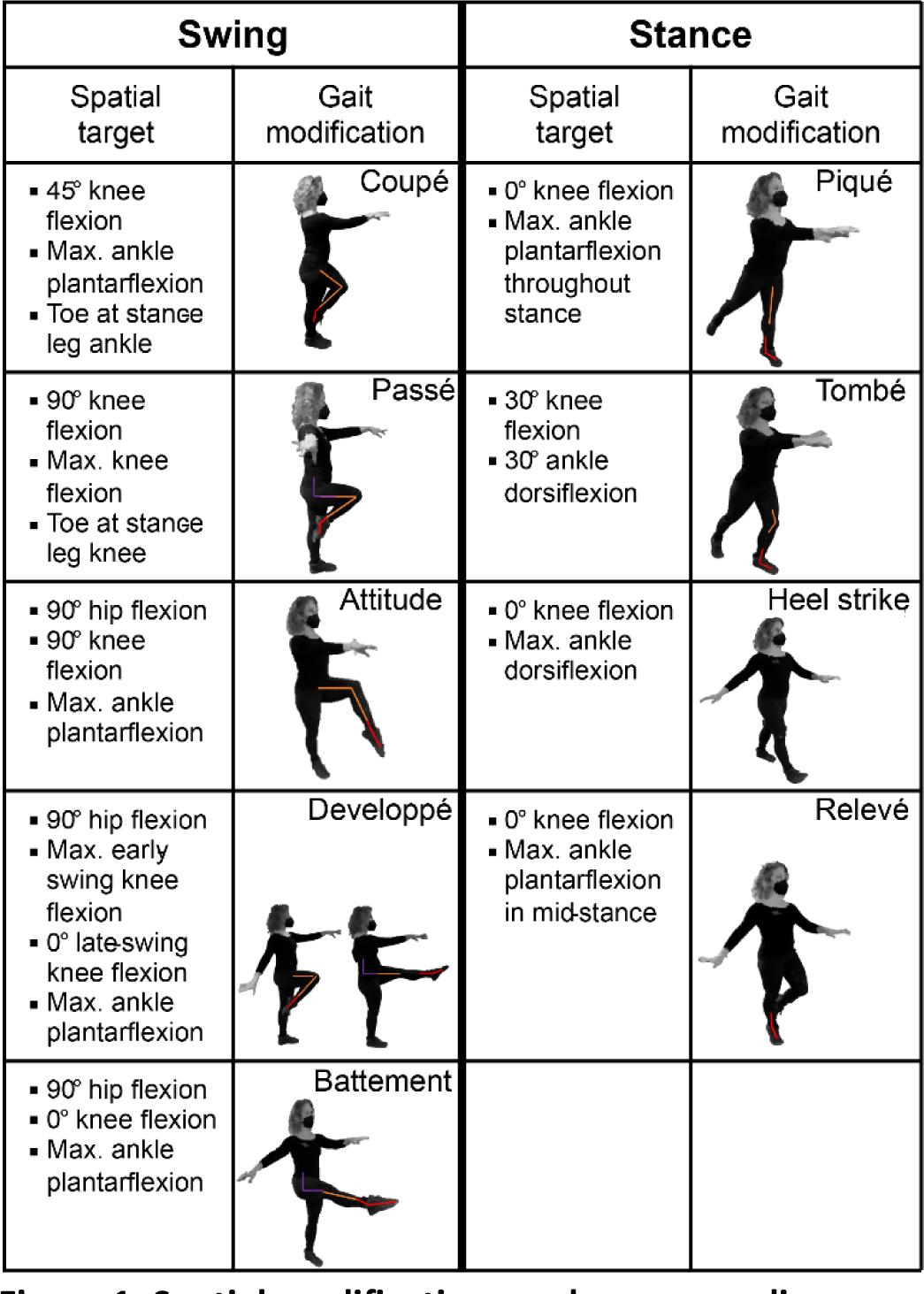
Spatial modifications and corresponding targets used in spatial and spatiotemporal rhythmic movement sequences (RMS). A) Spatial modifications. The left two columns correspond to modifications to swing-phase kinematics during movement, while the right two columns correspond to modifications to stance-phase kinematics. The bullets describe joint kinematics defining each spatial target variable for the corresponding modification. Deviations from these target values quantified RMS performance. The colored lines denote the hip (purple), knee (orange), and ankle (red) target values. *Adapted, with permission, from Rosenberg et al., 2023 [8]*.

Temporal RMS involved walking while performing repeating sequences of 2-6 steps of varying duration, with footfalls synchronized to musical pieces that served as external rhythmic cues [8]. Each temporal RMS consisted of only one temporal gait modification. The spatial aspects of movement were not constrained; participants could walk with any gait pattern and any speed that let them achieve the desired step sequence. Steps were performed at different rates relative to prescribed music: half steps spanned half of a beat (*&* in Figure 2), quick (q) steps spanned one beat (solid note in Figure 2), and slow (S) steps spanned two beats (open note in Figure 2). We defined three meter-based subclasses of temporal RMS from ballroom dance: *simple duple* (2-count), *complex duple* (2-count), and *waltz* (3-count) metric modifications, shown in Figure 2. Each RMS subclass contained three different step sequences, performed to the same rhythm and meter. For example, the *simple duple – qqSS* RMS involved taking two quick steps at a cadence of one step per beat, followed by two slow steps at a cadence of one step every two beats. Each modification class was expected to challenge different aspects of Western music listeners’ experiences [8, 14]. Participants performed duple and waltz modifications synchronously to modified versions of Libertango (by Astor Piazzolla, 1974) and Waltz No. 2 (by Dmitri Shostakovich, 1938), respectively, with superfluous accents and cues removed [8, 29]. All participants performed the duple modifications at 100 beats per minute (bpm). For the waltz modifications, the YA groups performed modifications to music at 80 bpm, while the music was slowed to 60 bpm for the OA and MCI groups to ensure that they could perform the modifications. Empirical tapping studies and theoretical models of motor-auditory synchronization suggest that the OA and MCI groups could be expected to exhibit better synchronization to the waltz 60bpm tempo than would the YA group to the 80bpm tempo [30-33]. However, we previously found that any differences in motor-auditory synchronization at 60bpm were small compared to within-group variability in RMS performance: the YA group did not perform waltz RMS less accurately than the OA group [8].

**Figure 2:**
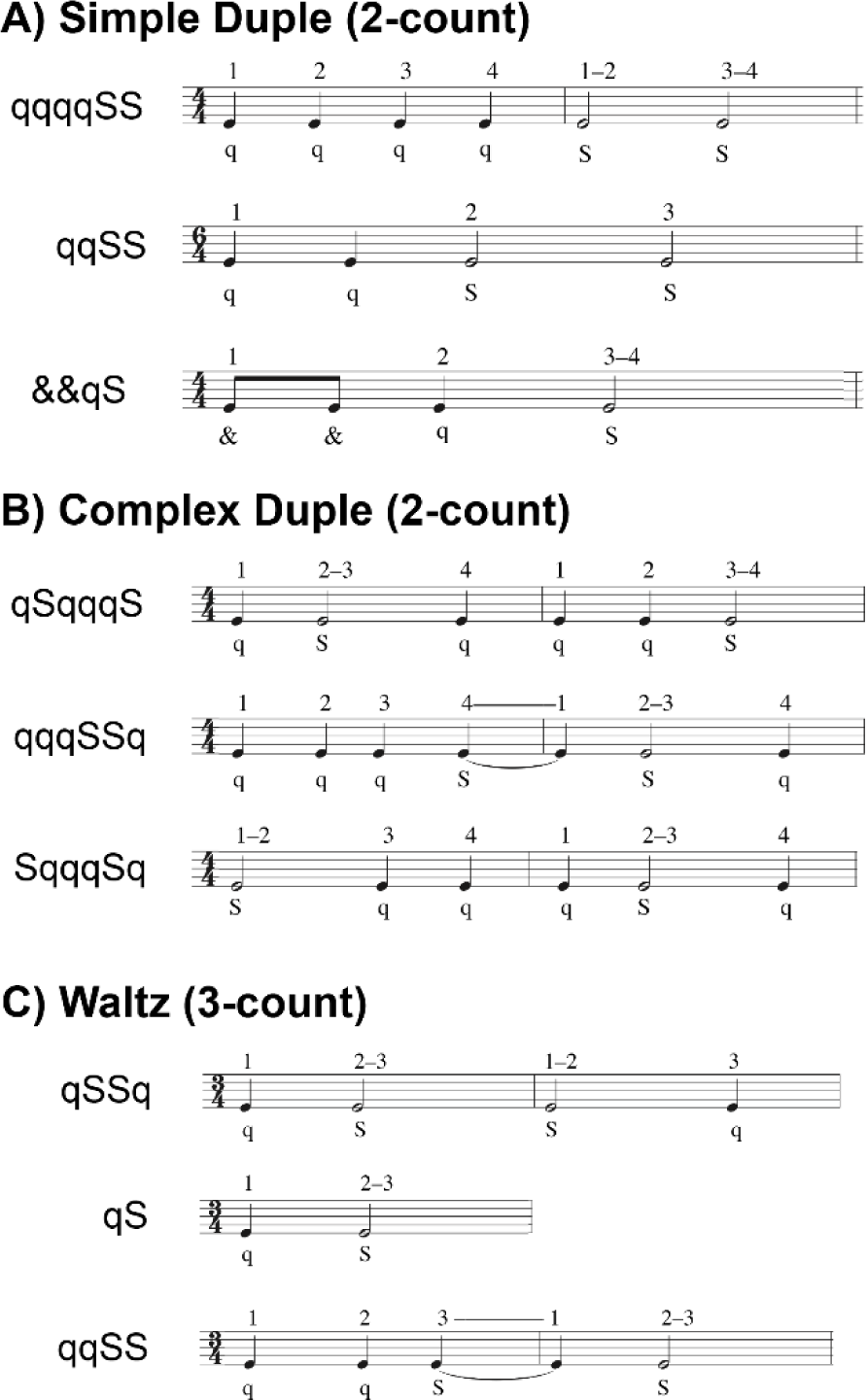
Rhythmic stepping sequences used in temporal and spatiotemporal rhythmic movement sequences (RMS). Each sequence consisted of 2-6 steps, synchronized to either a duple (2-count) or waltz (3-count) meter. Sequences were comprised of very-quick (*&* = half-beat per step), quick (*q* = one beat per step), and slow (*S* = two beats per step) steps. The numbers above each musical note reflect the beat count. A) Simple duple sequences were two-count rhythms with the strong beat on the downbeat and spanned 1-2 measures. B) Complex duple sequences were also two-count rhythms with a weak beat on the downbeat and spanned 2 measures. C) Waltz sequences were three-count rhythms spanning 1-2 measures. *Re-used, with permission, from Rosenberg et al., 2023 [8]*.

Each spatiotemporal RMS involved performing spatial and temporal modifications. Participants performed four spatiotemporal RMS consisting of different combinations of two spatial modifications and one temporal modification (Figure 1 & 2).

The RMS trial order was block-randomized for each participant. Spatial or temporal RMS were randomly selected to be performed first and second, followed by spatiotemporal RMS. Within each RMS class (spatial, temporal, spatiotemporal), the order of RMS sub-classes was randomized, as was the order of the three RMS within each subclass.

### RMS protocol

Participants performed each of the 22 RMS once. For each RMS, YA performed RMS while walking overground for four lengths of an 11-meter walkway. To mitigate the effects of fatigue on RMS performance, participants in the OA and MCI groups performed RMS for at least 11 meters (one walkway length) [8]. Participants who walked shorter distances also took more strides per walkway length. All participants performed each RMS for at least 15 strides. Participants could take as much time as needed to safely complete each trial. Across participants, trials typically lasted less than 180s, though longer trials took up to 360s. However, longer trials typically occurred once per participant, after which we reduced the number of walkway lengths to mitigate fatigue.

During RMS assessments, sagittal-plane hip, knee, and ankle kinematic timeseries were recorded at 128 Hz using Opal V2R inertial measurement units (APDM, Inc., Portland, USA). Fifteen sensors were attached to the forehead, sternum, lumbar region and bilaterally to the hands, wrists, upper arms, thighs, shanks, and feet in a standard configuration [34]. For each trial, joint kinematics were estimated using validated proprietary software (APDM Moveo Explorer) [35]. Computation of RMS performance is described in the next section.

Before performing each spatial RMS, participants watched a tutorial video of the sequence (two spatial modifications) being performed by an expert and received instructions on how to achieve the spatial targets of the movement (*i.e.*, discrete joint angle targets; Figure 1) [8]. Before performing each temporal RMS, participants watched a tutorial video that guided progressive entrainment of the trial’s rhythmic pattern in five steps of increasing complexity: clapping, tapping with one foot, shifting weight between the feet, marching in place, and walking to the rhythmic pattern. Our prior work found this approach to be an effective tool to teach new rhythms to dance-based therapy participants [17]. We found the technique useful for participants in the present study who had little music and dance experience. Before performing each spatiotemporal RMS, participants were reminded of the spatial targets (spatial) and step sequences (temporal) and could review the tutorial videos if needed.

Participants practiced each RMS with assessor feedback until the assessor determined that the participant understood the RMS. Practice typically spanned less than one walkway length. At the beginning of each walkway length during temporal trials, the assessor would clap the rhythm for two sequences to help participants identify the rhythm.

### RMS performance targets and quantification

Performance of each RMS was quantified as each participant’s ability to achieve pre-defined spatial targets (spatial and spatiotemporal RMS) and temporal targets (temporal and spatiotemporal RMS). Targets were specific to each modification and are described in Figure 1 & 2. Spatial targets were defined by sagittal-plane joint angles of the hip, knee, and/or ankle in the stance or swing phases for each RMS (Figure 1 & 3A). For example, the *Attitude* modification involved joint angle targets of 90-degree hip flexion, 90-degree knee flexion, and maximal ankle plantarflexion (Figure 3A, middle; stars denote target values).

**Figure 3:**
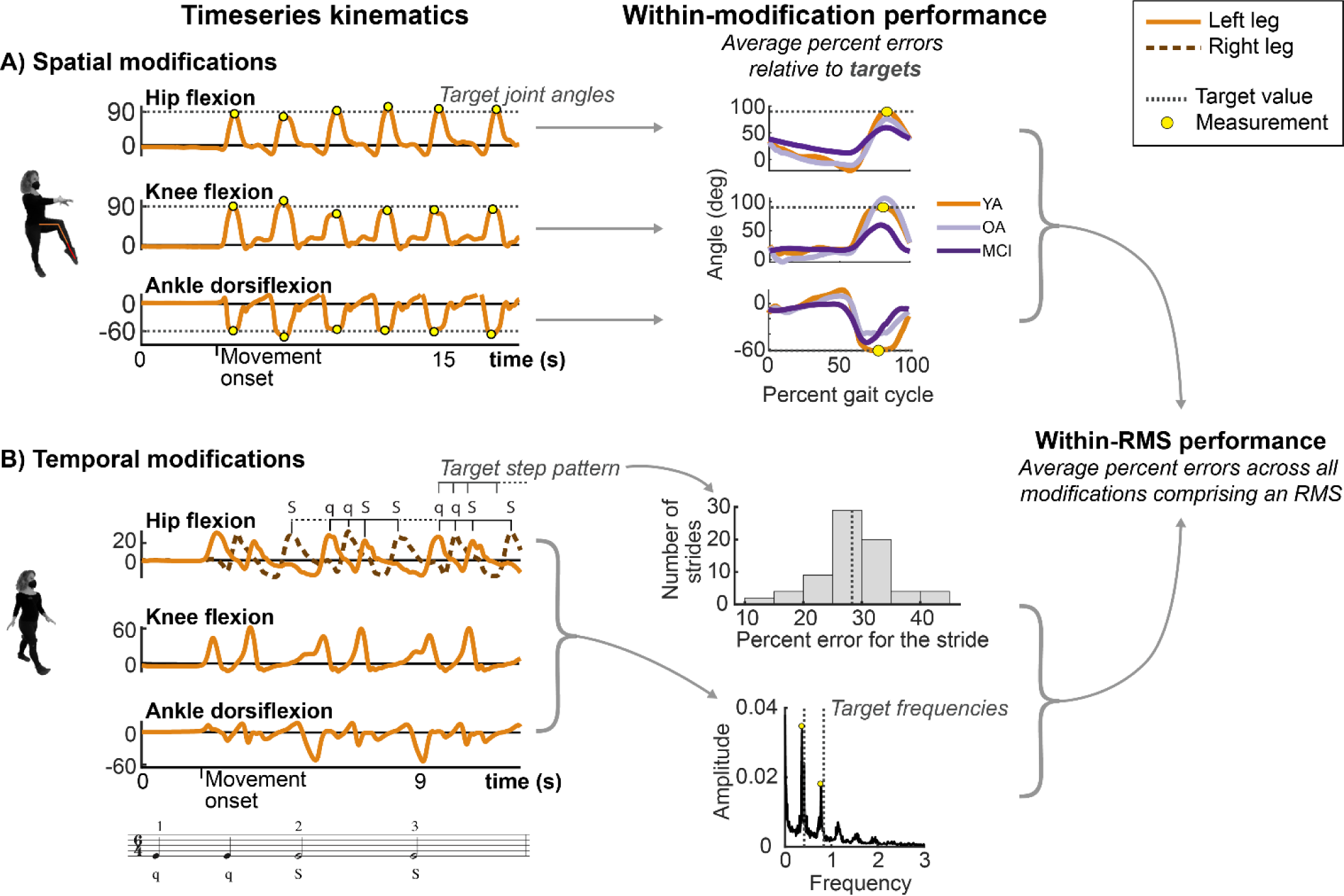
Calculation of error on spatial and temporal modifications comprising RMS. The left column shows timeseries data. The middle column depicts how percent errors computed within spatial and/ or temporal modifications comprising a given RMS. The right column depicts how percent errors within multiple modifications combine to produce error for each RMS. Dashed lines denote spatial and temporal targets. Yellow dots denote the portion of the stride (e.g., swing vs. stance) where the joint angles were compared to target values. A) One example spatial modification (*Attitude*). Percent errors of peak joint angles relative to target values were computed for each joint separately (left), then averaged across strides and target variables to compute and overall modification error (middle). The middle column shows stride-averaged data from one participant from each of the YA (orange), OA (gray), and MCI (purple) groups. B) One example temporal modification (bottom; *simple duple–quick-quick-Slow-Slow; qqSS*). The target rhythm is shown as musical notation below the temporal timeseries. Temporal targets were defined by step sequences, shown by the q-q-S-S notation. Deviations in the timing of kinematic peaks relative to the prescribed temporal pattern constituted error. The middle column shows a histogram of percent errors, which were averaged across strides (dashed line). The bottom middle plot depicts a Fast Fourier Transform of the timeseries data, in which the dominant frequencies (peaks) are compared to target frequencies (dashed lines). Temporal modification performance was defined as the average of these percent errors. Finally, to compute RMS performance error, all spatial and/or temporal modifications constituting an RMS were averaged (w*ithin-RMS performance*).

Temporal targets for each modification were defined by the modification’s pattern of quick (q) and slow (S) steps and the prescribed tempos (Figure 2). For example, the *simple duple – qqSS* modification (Figure 2A) involved performing two quick steps, followed by two slow steps at cadences of 100 steps per minute for the quick steps and 50 steps per minute for the slow steps.

For each RMS, performance was defined as the percent error relative to the spatial and temporal targets, averaged over all gait cycles, targets, and modifications comprising the RMS (Figure 3). For spatial modifications, percent errors were computed for executed joint angles relative to the prescribed joint targets evaluated at key instances of the gait cycle (stance or swing; Figure 3A, yellow dots). The percent errors were averaged across all strides and target joints for the modification (targets are specified in Figure 1; multiple strides are shown in Figure 3A, left). For temporal modifications, percent errors were computed for the temporal sequence of hip flexion angles relative to the prescribed step sequence (Figure 3B, timeseries peaks deviate from target sequence timing), and the dominant frequencies of the kinematics relative to target frequencies that would be expected by the pattern of quick and slow steps (Figure 3B, middle). This approach, rather than using step times, was necessary because foot contact events could not be reliably detected for all gait modifications using the inertial sensors. We identified dominant frequencies at the maxima of a Fast Fourier Transform of the joint kinematics (Figure 3B, middle). If a temporal modification consisted of quick and slow steps, we identified two dominant frequencies. If half, quick, and slow steps were included, we identified three dominant frequencies. For example, the *qqSS* simple duple sequence at 100 bpm would have target dominant frequencies of 100 (q) and 50 (S) bpm, or 1.67 and 0.83 steps/s.

To compute an overall percent error for each RMS, errors computed for each target and gait cycle in a modification were first computed for each variable of the modification’s target variables (e.g., yellow dots in Figure 3A and temporal error in Figure 3B). The errors within each modification were then averaged (Figure 3, middle). The RMS performance was then defined as the average percent error across modifications comprising that RMS (Figure 3, right). For example, RMS performance error on the spatiotemporal *coupé-*passé *– simple duple qqSS* was defined as the average of percent errors for the *coupé*, *passé*, and *simple duple qqSS* modifications. Finally, for RMS classes (spatial, temporal, spatiotemporal), percent errors for each RMS within the corresponding class were averaged. Lower spatial or temporal percent error implied better RMS performance.

### Music and dance relationships and proficiency assessments

To quantify individuals’ relationships to music, including their prior experiences and how they interact with music in daily life, we developed a Music Relationship Questionnaire (MRQ; Table 2) [36]. The MRQ was administered via REDCap and consisted of ten introspective questions on a Likert scale with seven response categories. A composite MRQ score was defined as the average score of the ten Likert scale questions, producing a maximum possible score of 7, with larger scores reflecting stronger relationships to music. The MRQ also included an open-ended question about active engagement in music and a multiple-choice selection of musical genre preferences (see *Supplemental – S3* for more information).

**Table 2:** List of questions asked for the Music Relationships Questionnaire (MRQ) and Dance Relationship Questionnaire (DRQ).

| Item | MRQ | DRQ | Scoring Scale** |
| --- | --- | --- | --- |
| 1 | Music is important in my life. | Dance is important in my life. | Strongly disagree = 1<br>Strongly agree = 7 |
| 2 | I listen to music in my typical day. | I dance in my typical day.* | Strongly disagree = 1<br>Strongly agree = 7 |
| 3 | I focus on what I am hearing while listening to music. | While listening to music, I often physically respond (ex: tapping, moving, clapping, nodding, snapping). | Strongly disagree = 1<br>Strongly agree = 7 |
| 4 | I focus on other things while music is playing. | I do not often move while music is playing. | Strongly agree = 1<br>Strongly disagree = 7 |
| 5 | I actively choose the music I listen to. | I actively choose to move when music is playing. | Strongly disagree = 1<br>Strongly agree = 7 |
| 6 | I usually listen to whatever music happens to be playing. | I usually dance to whatever music happens to be playing. | Strongly disagree = 1<br>Strongly agree = 7 |
| 7 | I play an instrument/sing. | I dance regularly. | Strongly disagree = 1<br>Strongly agree = 7 |
| 8 | I have played an instrument/sung for most of my life. | I have danced for most of my life. | Strongly disagree = 1<br>Strongly agree = 7 |
| 9 | I played an instrument/sang only as a child. | I danced only as a child. | Strongly disagree = 1<br>Strongly agree = 7 |
| 10 | In a typical day, I play an instrument/sing. | In a typical day, I dance.* | Strongly disagree = 1<br>Strongly agree = 7 |

\*Two items that were worded similarly and highly correlated (Pearson's $r = 0.89$ ) were averaged before computing the composite score for the DRQ.
\*\*The Scoring Scale was identical for the MRQ and DRQ. Note that the Scoring Scale on Item 4 is reversed relative to all other items.

Similarly, to quantify individuals’ relationships to dance, including their prior experience and how they interact with dance in daily life, we developed a Dance Relationship Questionnaire (DRQ; Table 2) [36]. The DRQ was administered via REDCap and consisted of ten introspective questions on a Likert scale with seven response categories. Like the MRQ, a composite DRQ score was initially defined as the average score of the 10 Likert scale questions, producing a maximum possible score of 7, with larger scores reflecting stronger relationships to dance. However, because questions 2 and 10 were similarly worded and responses were strongly correlated (Pearson’s *r* = 0.89), we took the average of these questions before computing the composite score across all questions. The DRQ also included an open-ended question about active engagement in dance and a multiple-choice selection of dance style preferences (see *Supplemental – S3* for more information).

To quantify individuals’ proficiency in perceiving rhythms and executing motor commands in-sync with those rhythms, we developed an objective rhythm assessment (RA; Table 3) [36]. The RA was conducted in-person or virtually by trained personnel and took 10-15 minutes to complete. The assessment consisted of three parts: 1) To assess participants’ abilities to perceive, comprehend, and reproduce rhythmic patterns from *auditory* stimuli, participants listened to four recordings of rhythmic patterns of quick and slow claps (listened three times per pattern), then attempted to accurately clap each rhythm twice (Table 3; items 1-4). These tasks are denoted *auditory rhythm reproduction* tasks. 2) To assess participants’ abilities to perceive, comprehend, and replicate rhythmic patterns from *visual* stimuli, participants read two measures of Western music notation in 4/4 time and attempted to accurately clapback the rhythm (Table 3; item 5). This task is denoted *visual rhythm reproduction* tasks. Participants could practice the rhythms for up to one minute before their clapping was scored and were instructed to attempt to clap the rhythm even if they could not read musical notation. Anecdotally, participants who verbally indicated they could not read musical notation typically received a score of zero on this item. The audio of all clapped rhythms was recorded for scoring. 3) To assess participants’ abilities to recognize different meters in music, participants listened to five music passages, then identified whether that passage consisted of either “two’s” (*duple*), “three’s” (*triple*), or “other” meters (Table 3; items 6-10). These tasks are denoted *auditory meter recognition* tasks. All passages were less than 60 seconds in length and played up to three times for the participant. The RA composite score was defined as the sum of scores on the 10 items, with a maximum score of 10 points (Table 3). Correct answers received one point, partially correct answers (e.g., correct on one of two clapbacks on the *auditory* or *visual rhythm production* tasks) received one-half point, and incorrect answered received zero points. RA instructions are provided in *Supplemental – S2*.

**Table 3:** Description of assessment items in the Rhythm Assessment (RA).

| Item | Item Description | Scoring Scale – Response (Points) |  |  |
| --- | --- | --- | --- | --- |
| 1 | Auditory rhythm reproduction A | Correct (1) | Partial (0.5) | Incorrect (0) |
| 2 | Auditory rhythm reproduction B |  |  |  |
| 3 | Auditory rhythm reproduction C |  |  |  |
| 4 | Auditory rhythm reproduction D |  |  |  |
| 5 | Visual rhythm reproduction A |  |  |  |
| 6 | Auditory meter recognition A (Duple) | Duple (1) | Waltz (0) | Other (0) |
| 7 | Auditory meter recognition B (Waltz) | Duple (0) | Waltz (1) | Other (0) |
| 8 | Auditory meter recognition C (Duple) | Duple (1) | Waltz (0) | Other (0) |
| 9 | Auditory meter recognition D (Waltz) | Duple (0) | Waltz (1) | Other (0) |
| 10 | Auditory meter recognition E (Other) | Duple (0) | Waltz (0) | Other (1) |

“Partially correct” indicated that the participant clapped the rhythm correctly once and incorrectly once.
*Auditory* items involved listening to a rhythm, then clapping it back.
The *Visual* item involved reading a sequence of 10 musical notes (eighth, quarter, half beats) then clapping the notes.
*Meter recognition* items involved listening to a rhythm, then identifying whether the meter was “twos” (two-count), “threes” (three-count), or “other.”

The RA documentation and audio recordings, along with the data used in this manuscript and a data dictionary, are freely available in the supplemental materials (see *Supplemental – S1* for a description of these files).

### Statistical analysis

To determine if groups differed in their experiences with dance and music, as well as their rhythmic proficiency, we tested for differences in MRQ, DRQ, and RA scores between all pairs of groups (YA, OA, and MCI) using independent-sample t-tests (α = 0.05). Because our sample was larger than that in [8] and [36], we replicated the study’s group-wise comparisons of spatial, temporal, and spatiotemporal RMS performance using independent-sample t-tests (α = 0.05). Because we were previously interested in how age and cognitive status impacted RMS performance, we only compared RMS performance between YA and OA (age effect) and between OA and MCI (cognitive status effect) [8].

To characterize relationships between the MRQ, DRQ, and RA, we performed univariate linear regression of the composite scores between pairs of the MRQ, DRQ, and RA. Similarly, to determine if the MRQ, DRQ, and RA were associated with measures of cognitive and motor-cognitive function, we performed univariate linear regression between these composite scores and scores on the MoCA, Trail Making Task (B-A), Reverse Corsi Blocks test, and the Body Position Spatial Task. Regressions were performed separately for each assessment pair and separately for the YA, OA, and MCI groups.

To determine if, within each participant group, stronger relationships to music (higher MRQ composite scores) were associated with better temporal modulation ability (lower RMS performance error), we performed linear regression between MRQ composite scores and temporal RMS performance separately for each group. Analyzing groups separately was based on our prior finding that RMS performance differed between groups [8]. Similarly, to determine if, within groups, stronger relationships to music were associated with better spatiotemporal modulation ability, we performed linear regression between MRQ composite scores and spatiotemporal RMS performance.

To determine if, within groups, stronger relationships to dance (higher DRQ composite scores) were associated with better spatial modulation ability, we performed linear regression between DRQ composite scores and spatial RMS performance. Similarly, to determine if, within groups, stronger relationships to dance were associated with better spatiotemporal RMS performance, we performed linear regression between DRQ scores and spatiotemporal RMS performance.

To determine if stronger rhythmic proficiency (higher RA composite scores) was associated with better temporal and spatiotemporal modulation ability, we performed linear regression between RA composite scores and temporal and spatiotemporal RMS performance, respectively, for each group separately.

Finally, because we identified associations between RA composite scores and RMS performance, we sought to determine if the ability to reproduce auditory rhythms and identify meter in musical pieces differentially impacted RMS performance. We conducted an exploratory regression analysis of the *auditory rhythm reproduction* and *auditory meter recognition* sub-components of the RA against temporal and spatiotemporal RMS separately for each group.

For all analyses, we report regression accuracy as coefficients of determination (r^2^), regression slopes, and significance levels according to Wald tests (α = 0.05). Group-level comparisons of demographics and clinical characteristics were performed using the software package *R*. Regression analyses were performed using MATLAB 2021b (Mathworks, Natick, USA).

## Results

*Clinical assessments suggest worse cognitive function in adults with MCI, compared to older adults* Clinical assessments revealed group-level differences in motor-cognitive and cognitive function (Table 1). Compared to the OA group, the YA group exhibited better performance on assessments of working memory (Reverse Corsi Blocks; p < 0.001) and motor-cognitive integration (Body Position Spatial Task; p < 0.001). Similarly, the MCI group exhibited worse performance than the OA group on assessments of global cognitive function (MoCA; p = 0.005), working memory (Reverse Corsi Blocks; p = 0.035), and set-shifting (Trail Making Test; p = 0.050).

### Music relationships, rhythmic proficiency, and RMS performance differed between groups

Relationships to music and rhythmic proficiency differed between groups (Figure 4A). The YA and OA groups exhibited stronger relationships to music and greater rhythmic proficiency than the MCI group.

**Figure 4:**
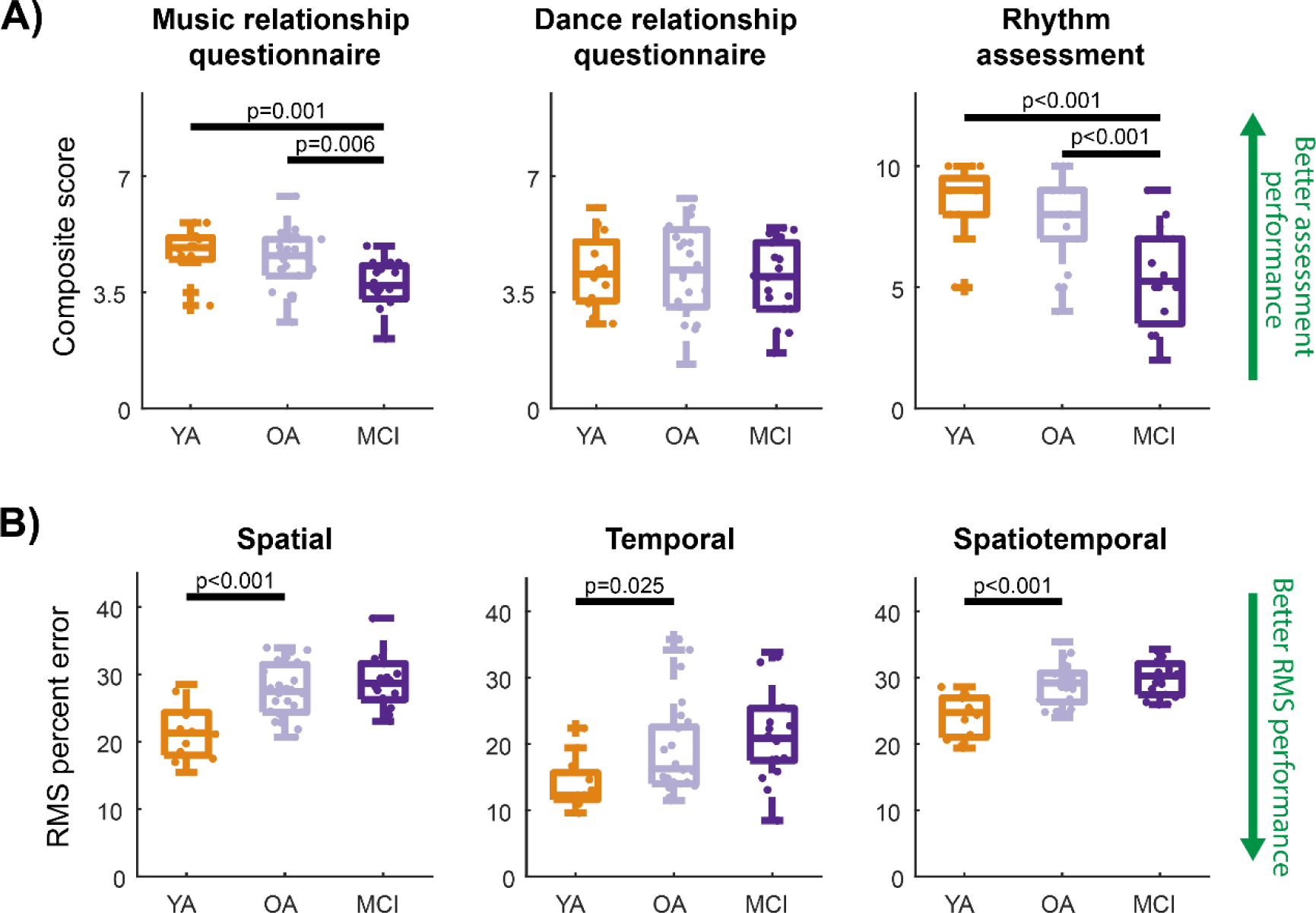
Boxplots representing distributions of Music Relationship Questionnaire (MRQ), Dance Relationship Questionnaire (DRQ), and Rhythm Assessment (RA) scores. A) MRQ, DRQ, and RA scores for the three participant groups (YA, OA, MCI). Higher MRQ and DRQ scores (max = 7) reflect stronger music and dance relationships, respectively. Higher RA scores (max = 10) reflect greater rhythmic proficiency. B) RMS performance error on each of spatial, temporal, and spatiotemporal RMS for the three participant groups. For both plots, dots represent individual participants. Higher composite scores indicate better performance on the MRQ, DRQ, and RA (upper green arrow). Lower RMS error indicates better performance on RMS (lower green arrow). Each spatial and temporal RMS, error was averaged across 9 RMS from their respective domains and spatiotemporal error was averaged across 4 RMS. For all boxplots, p-values denote significant differences according to independent-samples t-tests (α = 0.05).

On the MRQ, the YA group scored an average of 1.0 points (out of 7) higher (p = 0.001) and the OA group scored 0.7 points higher (p = 0.006) than the MCI group. The YA and OA groups exhibited similar relationships to music, as indicated by similar MRQ scores (p = 0.228). On the RA, the YA group scored an average of 3.2 points (out of 10) higher (p < 0.001) and the OA group scored 2.4 points higher (p < 0.001) compared to the MCI group. YA and OA groups exhibited similar levels of rhythmic proficiency, as indicated by similar RA scores (p = 0.223). No groups exhibited differences in the strength of their dance relationships, as indicated by similar DRQ scores (p > 0.575).

Compared to OA, the YA group exhibited better RMS performance, as indicated by lower percent error on spatial (p < 0.001), temporal (p = 0.025), and spatiotemporal (p < 0.001) RMS (Figure 4B). The OA group did not perform RMS with different accuracy than the MCI group (all p > 0.257).

### MRQ, DRQ, and RA composite scores were not strongly correlated across groups

Greater MRQ composite scores were associated with greater DRQ and RA composite scores only in the OA group (r^2^ < 0.27; p < 0.024; Table 4). Conversely, greater DRQ composite scores were associated with greater RA scores only in the YA group (r^2^ < 0.43; p < 0.021).

**Table 4:** Univariate linear regression results testing for covariation between MRQ, DRQ, RA composite scores.

| Questionnaire | Questionnaire / assessment | YA |  |  | OA |  |  | MCI |  |  |
| --- | --- | --- | --- | --- | --- | --- | --- | --- | --- | --- |
| | | $r^2$ | slope | p | $r^2$ | slope | p | $r^2$ | slope | p |
| MRQ | DRQ | 0.28 | 0.8 | 0.080 | 0.20 | 0.8 | **0.024** | 0.04 | 0.4 | 0.399 |
|  | RA | 0.26 | 0.3 | 0.087 | 0.27 | 0.2 | **0.010** | 0.22 | 0.2 | 0.064 |
| DRQ | RA | 0.43 | 0.5 | **0.021** | 0.00 | 0.0 | 0.848 | 0.17 | 0.2 | 0.109 |

Abbreviations: YA, younger adults; OA, older adults without mild cognitive impairment; MCI, older adults with mild cognitive impairment; MRQ, Music Relationship Questionnaire; DRQ, Dance Relationship Questionnaire; RA, Rhythm Assessment
For each group, regression results are shown using the coefficient of determination ( $r^2$ ), regression slope, and p-value ( $p$ ) denoting a slope that is significantly different from zero ( $\alpha = 0.05$ ). Bold p-values denote $p < 0.05$ .

### Clinical measures of cognitive and motor-cognitive function were not associated with the MRQ, DRQ, or RA scores

Across four clinical measures of cognitive and motor-cognitive function, we identified only one significant association between the Body Position Spatial Task and MRQ in the OA group (r^2^ = 0.37; slope = 0.06; p = 0.001; Table 5). Scores on the MoCA, Trail Making Task (B-A), and Reverse Corsi Blocks test were not associated with the MRQ, DRQ, or RA in any group.

**Table 5:** Univariate linear regression results between clinical assessments of cognitive and motor-cognitive function and the MRQ, DRQ, and RA composite scores.

| Questionnaire / assessment | Assessment | YA | OA | MCI |
| --- | --- | --- | --- | --- |
| MRQ | MoCA | 0.00 | 0.00 | 0.12 |
|  | Trails B-A | 0.02 | 0.00 | 0.02 |
|  | Reverse Corsi blocks | 0.06 | 0.05 | 0.04 |
|  | Body position spatial task | 0.00 | **0.37** | 0.06 |
| DRQ | MoCA | 0.01 | 0.01 | 0.00 |
|  | Trails B-A | 0.00 | 0.00 | 0.09 |
|  | Reverse Corsi blocks | 0.00 | 0.00 | 0.01 |
|  | Body position spatial task | 0.14 | 0.01 | 0.22 |
| RA | MoCA | 0.01 | 0.01 | 0.00 |
|  | Trails B-A | 0.04 | 0.01 | 0.04 |
|  | Reverse Corsi blocks | 0.13 | 0.03 | 0.19 |
|  | Body position spatial task | 0.14 | 0.11 | 0.03 |

Abbreviations: YA, younger adults; OA, older adults without mild cognitive impairment; MCI, older adults with mild cognitive impairment; RMS, Rhythmic Movement Sequence; MRQ, Music Relationship Questionnaire; DRQ, Dance Relationship Questionnaire; RA, Rhythm Assessment; MoCA, Montreal Cognitive Assessment; Trails B-A, Trail Making Task items B-A.
For each group, regression results are shown using the coefficient of determination ( $r^2$ ), boldfaced values denote a correlation with slope that was significantly different from zero ( $\alpha = 0.05$ ) without correction for multiple comparisons.

### Higher MRQ scores were associated with reduced temporal RMS errors in young and older adults without MCI

In participants without MCI, individuals with higher MRQ scores (stronger music relationships) exhibited lower error on temporal RMS (i.e., better temporal RMS performance). MRQ scores were associated with temporal RMS errors in the YA (r^2^ = 0.41; slope = -3.2; p = 0.026; Table 6; orange in Figure 5A, top) and OA groups (r^2^ = 0.18; slope = -3.5; p = 0.030; grey in Figure 5A, top), explaining a small-to-moderate amount of the variance in temporal RMS errors. Conversely, the MCI group did not exhibit significant associations between MRQ scores and temporal RMS performance (r^2^ = 0.06; p = 0.314; purple in Figure 5A, top). No group exhibited significant associations between MRQ scores and spatiotemporal RMS performance (all r^2^ < 0.15; p > 0.114; Figure 5A, bottom).

**Figure 5:**
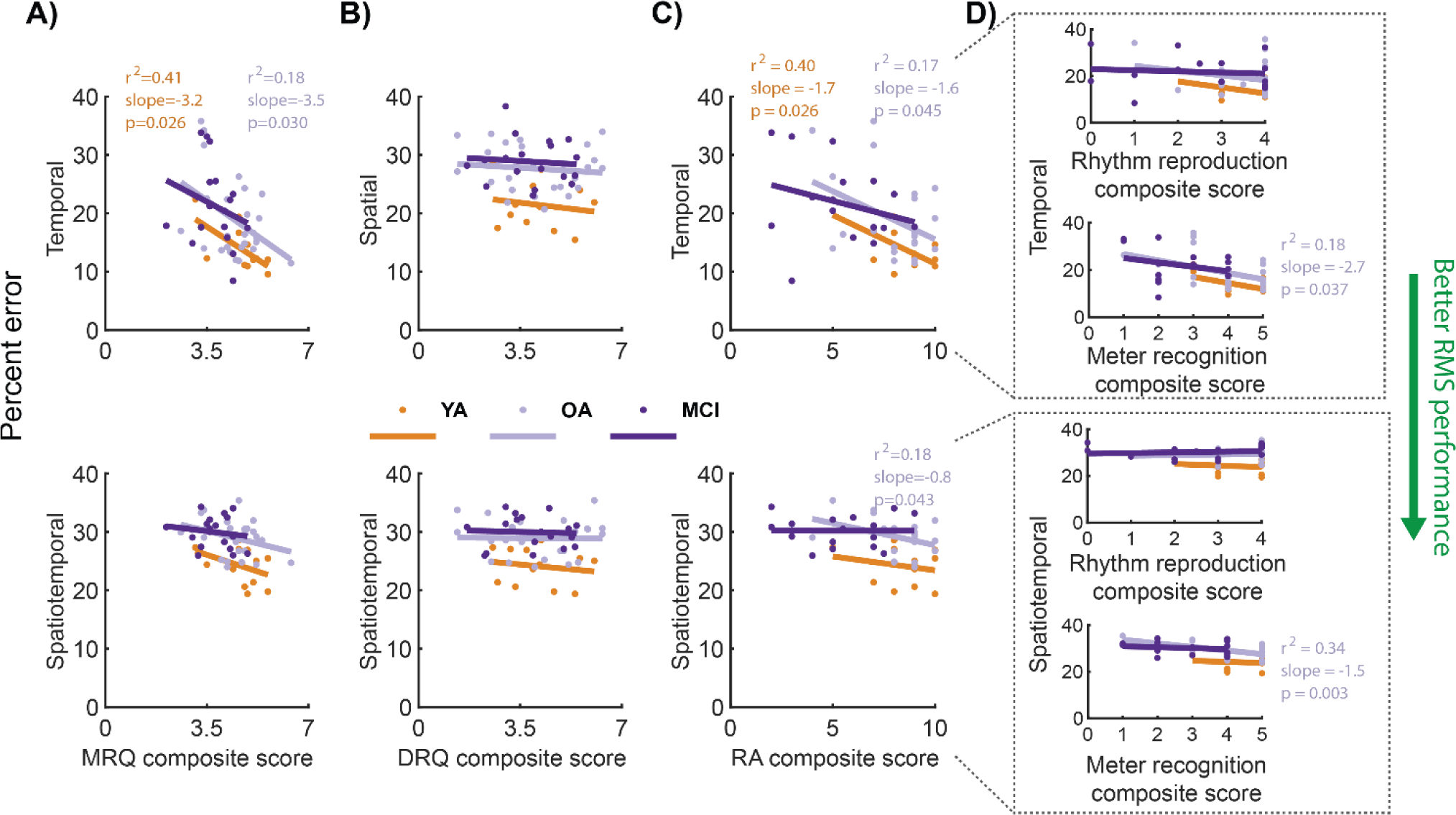
Linear regression testing for within-group associations between RMS performance and each of the MRQ, DRQ, and RA. Each dot represents a single participant. Lines indicate within-group linear fits. Colors denote the groups (YA: orange, OA: gray, MCI: purple). Regression R-squared values, slopes, and p-values are shown on the corresponding plots for fits that were significantly different from zero (Wald tests; *α* = 0.05), with colors corresponding to groups. Each of the spatial and temporal RMS percent errors were averaged across the 9 respective RMS and spatiotemporal errors were averaged across 4 RMS. Lower RMS error indicates better performance (green arrow), on average, across RMS within the corresponding RMS class. A) MRQ vs. temporal (top) and spatiotemporal (bottom) RMS performance errors. Higher MRQ scores represent stronger music relationships. B) DRQ vs. spatial (top) and spatiotemporal (bottom) RMS performance errors. Higher DRQ scores represent stronger dance relationships. C) Comparisons of RA with percent errors on temporal (top) and spatiotemporal (bottom) RMS. Higher RA scores imply greater rhythmic proficiency. D) Comparisons of RA subcomponents—*auditory rhythm reproduction* (max score = 4) and *auditory meter recognition* (max score = 5)—to temporal (top) and spatiotemporal (bottom) RMS.

**Table 6:** Univariate linear regression results between MRQ, DRQ, and RA composite scores and RMS performance.

| Questionnaire / assessment | RMS class | YA |  |  | OA |  |  | MCI |  |  |
| --- | --- | --- | --- | --- | --- | --- | --- | --- | --- | --- |
| | | $r^2$ | slope | p | $r^2$ | slope | p | $r^2$ | slope | p |
| MRQ | Temporal | 0.41 | -3.21 | **0.026** | 0.18 | -3.49 | **0.030** | 0.06 | -2.62 | 0.317 |
|  | Spatiotemporal | 0.15 | -1.63 | 0.215 | 0.10 | -1.23 | 0.114 | 0.02 | -0.62 | 0.551 |
| DRQ | Spatial | 0.03 | -0.60 | 0.610 | 0.01 | -0.30 | 0.575 | 0.01 | -0.29 | 0.732 |
|  | Spatiotemporal | 0.03 | -0.47 | 0.612 | 0.00 | -0.03 | 0.942 | 0.00 | -0.12 | 0.841 |
| RA | Temporal | 0.40 | -1.67 | **0.026** | 0.17 | -1.65 | **0.045** | 0.08 | -0.91 | 0.299 |
|  | Spatiotemporal | 0.05 | -0.48 | 0.498 | 0.18 | -0.75 | **0.043** | 0.00 | 0.00 | 0.994 |
| RA rhythm reproduction | Temporal | 0.21 | -2.6 | 0.135 | 0.05 | -2.0 | 0.296 | 0.01 | -0.5 | 0.722 |
|  | Spatiotemporal | 0.02 | -0.7 | 0.667 | 0.00 | 0.2 | 0.791 | 0.02 | 0.3 | 0.623 |
| RA meter recognition | Temporal | 0.26 | -2.6 | 0.090 | 0.18 | -2.7 | **0.037** | 0.09 | -1.9 | 0.265 |
|  | Spatiotemporal | 0.01 | -0.4 | 0.743 | 0.34 | -1.5 | **0.003** | 0.03 | -0.4 | 0.534 |

Abbreviations: YA, younger adults; OA, older adults without mild cognitive impairment; MCI, older adults with mild cognitive impairment; RMS, Rhythmic Movement Sequence; MRQ, Music Relationship Questionnaire; DRQ, Dance Relationship Questionnaire; RA, Rhythm Assessment For each group, regression results are shown using the coefficient of determination ( $r^2$ ), regression slope, and p-value ( $p$ ) denoting a slope that is significantly different from zero ( $\alpha = 0.05$ ). Bold p-values denote $p < 0.05$ .

### DRQ scores were not associated with spatial or spatiotemporal RMS performance

No group exhibited significant associations between DRQ scores and spatial or spatiotemporal RMS performance (all r^2^ < 0.03; p > 0.575; Table 6; Figure 5B).

### Higher RA scores were associated with reduced temporal RMS errors in young and older adults without MCI

In participants without MCI, individuals with higher RA scores tended to exhibit better temporal RMS performance (i.e., lower error). Negative associations between RA scores and temporal RMS error were observed in the YA (r^2^ = 0.40; slope = -1.7; p = 0.026; Table 6; orange in Figure 5C, top) and OA groups (r^2^ = 0.22; slope = -1.7; p = 0.021; grey in Figure 5C, top), explaining a small-to-moderate amount of variance in temporal RMS errors. Conversely, the MCI group did not exhibit significant associations between RA scores and temporal RMS performance, in part due to heterogeneity in RMS performance across individuals with low RA scores (r^2^ = 0.11; p = 0.210; purple in Figure 5C, top). Only the OA group exhibited an association between better RA scores and better spatiotemporal RMS performance (all r^2^ = 0.18; slope = -0.75; p > 0.043; Figure 5C, bottom).

### Auditory rhythm reproduction and auditory meter recognition scores were only associated with RMS performance in older adults

When analyzing the RA sub-components, *auditory rhythm reproduction* (4 items in the RA) alone was not associated with temporal or spatiotemporal RMS performance in any group (r^2^ < 0.21; p > 0.135; Table 6; Figure 5D). For the second RA sub-component, better *auditory meter recognition* (5 items in the RA), was associated with better temporal (r^2^ > 0.18; slope = -2.7; p = 0.037) and spatiotemporal (r^2^ > 0.34; slope = -1.5; p = 0.003) RMS only in the OA group (Figure 5D).

## Discussion

This study shows that in young and older adults without MCI, stronger relationships to music and better rhythmic proficiency are associated with a better ability to accurately modulate temporal aspects of movement during RMS. Conversely, these relationships are not apparent in adults with MCI, suggesting that cognitive deficits associated with MCI hinder the ability to transform relationships to music or rhythmic proficiency into an ability to modulate temporal aspects of movement during RMS. Our central hypothesis was, therefore, only supported for temporal RMS in younger and older adults without MCI. Stronger relationships to dance do not appear to impact the ability to perform RMS involving spatial modifications to movement in any group. These findings suggest that only in the absence of MCI-related cognitive deficits will individuals’ relationships to music or rhythmic proficiency be indicative of the complexity of musical rhythms and step sequences that they can successfully perform during dance-based therapies [3, 5, 6].

Stronger relationships to music and rhythmic proficiency indicate a better ability to accurately modulate temporal features of movement during RMS in adults without MCI, but not with MCI. In adults without MCI, these associations suggest that constructs that are developed by prior relationships to music and support rhythmic proficiency also improve the ability to perceive rhythm and meter and entrain movement to these rhythms. For example, individuals with stronger music relationships may exhibit more precise mapping of temporal stimuli between auditory processing and motor planning regions in the brain [14, 32]. Such mapping may enable more accurate entrainment of step timing to music-based auditory cues during temporal RMS [37]. Given the likelihood of similar perception-action pathways invoked by the RA and temporal and spatiotemporal RMS [31, 32], it is reasonable to expect that within-group differences in RA and RMS performance are driven by similar constructs, which merits future investigation. For example, the ability to identify meter in musical pieces appears to be moderately predictive of temporal RMS performance in the absence of MCI. Our findings suggest that this skill may be developed through prior experiences with either music or dance.

Cognitive deficits in adults with MCI may have masked associations between music relationships or rhythmic proficiency and temporal RMS performance. Individual-specific working memory deficits in the MCI group may explain the more variable temporal RMS performance and may be exacerbated for longer temporal sequences. For example, we previously found that adults with MCI performed longer temporal RMS sequences (6-step duple sequences) less accurately than older adults and that lower temporal RMS performance was associated with worse working memory (i.e., lower scores on the Reverse Corsi Blocks test) [8, 24]. Note that the group differences in RMS performance presented in the present study are consistent with those previously reported using a subset of participants included here [8]. Similarly, lower rhythmic proficiency in adults with MCI, compared to those without MCI, suggests that attentional or motor-cognitive deficits accompanying MCI diagnosis may manifest as a reduced ability to perceive musical cues and entrain motor commands to music [14]. Despite adults with MCI reporting weaker relationships to music than older adults without MCI, our findings suggest individual differences in music and dance relationships are not driven by declines in cognitive or motor-cognitive function. Further research is required to determine the extent to which MCI-related impairments contribute to declines in rhythmic proficiency, which may impact individuals’ abilities to synchronize movements to auditory cues during dance-based therapies [12-14].

Unlike music relationships, stronger relationships to dance are not indicative of a better ability to accurately perform spatial RMS. Rather, our prior study found that differences in spatial RMS performance are better explained by differences in age-related motor and cognitive function [8]. Therefore, stronger dance relationships do not likely enhance constructs that are beneficial to spatial RMS performance consistently across individuals. For example, the experiences assessed by the DRQ neither require nor develop dynamic balance or the large joint ranges of motion required by spatial RMS [38]. Prior experience in non-dance activities like gymnastics may, therefore, be a better indicator of an individual’s ability to perform spatial RMS [39, 40]. Identifying specific subcomponents of dance relationships or other dance-related experiences that predict the ability to modulate spatial aspects of movement is an interesting future research direction. For example, only three participants reported a preference for ballet, which was the basis for the spatial modifications during RMS (*Supplemental – S3*). Adjusting the modifications comprising RMS to be derived from dance styles that more-strongly align with participants’ preferences may yield stronger relationships between DRQ scores and spatial RMS performance.

Better rhythmic proficiency, but not relationships to music or dance, may be indicative of better spatiotemporal RMS performance. However, we observed this trend only in older adults without MCI. As discussed in the prior paragraph, constructs not evaluated by the MRQ, DRQ, or RA likely impact spatial RMS performance, which constitutes half of the spatiotemporal RMS error. The relationship between better rhythmic proficiency and temporal RMS performance likely also drives better spatiotemporal RMS performance. The lack of association in young adults is not necessarily surprising; participants may choose to prioritize spatial, rather than temporal aspects of movement, potentially decoupling rhythmic proficiency from spatiotemporal RMS performance. However, our prior findings suggest that differences in motor function (e.g., balance) or cognitive function (e.g., set shifting), appear to be stronger predictors of the ability to perform spatiotemporal RMS [8, 23, 24, 41].

Several factors limit the generalizability and interpretation of our results. Limitations related to our experimental protocol and RMS performance quantification are discussed in [8]. While the OA and MCI group sample sizes in this study were larger than in [8] and [36], still larger sample sizes would improve confidence in our regression results for groups with highly variable RMS performance. Further, our analyses did not adjust for within-group effects of motor or cognitive function, or their interactions with music or dance relationships, on RMS performance [8]. Complex interactions between motor and cognitive decline, music or dance relationships, and rhythmic proficiency in individuals with MCI, may contribute to RMS performance. Future larger-sample studies could use multivariate regression techniques to extend our understanding of how individual differences in motor and cognitive function moderate the impacts of music and dance relationships or rhythmic proficiency on RMS performance.

Kinematic estimates from the APDM Mobility Lab software may also induce errors, particularly at large joint excursions for which the software and inertial sensors have not been validated [34, 35, 42, 43]. Prior studies of large joint ranges of motion suggest that IMU-based kinematic errors can reach 5-8 degrees, which may introduce non-trivial errors in RMS performance [44, 45]. However, these errors would likely reduce our ability to detect inter-individual differences in RMS performance: extreme joint angles would likely increase RMS error for the best performers. Further, experiment administrators reported, anecdotally, that participants with worse RMS percent errors struggled more to perform RMS.

Additionally, the MRQ, DRQ, and RA represent a preliminary characterization of music relationships, dance relationships, and rhythmic proficiency, respectively. The questionnaire/assessment items encompassed multiple constructs that may impact RMS performance. Computing MRQ, DRQ, and RA composite scores by averaging the scores for all items may mask more nuanced associations between music and dance relationships and RMS performance. Further, moderate covariation between the MRQ, DRQ, and RA scores suggests that these assessments encode some overlapping constructs. Analyzing individual questionnaire items may reveal subcomponents of music and dance relationships that are more strongly associated with RMS performance. Further, established questionnaires, such as the Goldsmith’s Music and Dance Sophistication Indices and the Barcelona Music Reward Questionnaire test similar perceptual and motor constructs (e.g., active listening, singing ability, sensory-motor) as those assayed in the MRQ, DRQ, and RA [46-48]. These previously established questionnaires include a broader range of questions, from proficiency to emotional engagement. The MRQ and DRQ were developed to assess broad relationships with music and dance, and did not include more nuanced emotional or social concepts. Unlike the MRQ, the DRQ assessed active, but not passive aspects of dance relationships. As dance observation is known to impact neural activity and dance performance, recording information about dance relationships will be valuable for future studies [27, 47, 49]. The inclusion of the additional constructs described above could provide a more detailed understanding of how prior experience with music or dance impacts RMS performance. Including a broader set of questions from the MRQ, DRQ, and RA, as well as established questionnaires, may be useful to identify specific aspects of music and dance relationships that impact the ability to perform dance-like movements.

Further, we did not control for individual preferences for different musical genres or dance styles, active (e.g., singing, playing an instrument, dancing) versus passive (e.g., listening to music, watching dance) engagement in music or dance, and extent of experience, all of which may affect the relationships between music or dance relationships and RMS performance. All participants in the present study reported listening to one or more Western musical genres. However, non-Western music listeners may struggle to identify rhythm and meter in Western musical pieces, possibly reducing RMS performance [14]. Most participants in the present study reported listening to classical music, and classical music listeners tended to perform better on temporal RMS, which used classical music pieces (see *Supplemental – S3* for more information). However, this trend may not be specific to classical music.

Similarly, we did not find that participants who reported active music or dance engagement performed better on RMS than those who did not (see *Supplemental – S3* for more information). Finally, we did not directly measure the extent (e.g., number of years) of experience in music or dance, which may impact RMS performance. Characterizing how group or individual differences in RMS performance generalize across preference, engagement style, and extent of experience in music and dance would improve our understanding of the clinical utility of RMS.

Finally, future studies should characterize the extent to which RMS performance is associated with participants’ capabilities within the many other dance styles and musical genres encountered during dance-based therapies, as well as the participant-reported difficulty, efficacy, and engagement in such therapies. Determining the extent to which RMS, which incorporate ballet fundamentals with classical musical selections, generalize to other music or dance styles would support the utility of RMS as a tool to quantify and predict therapeutic performance. Spatial and temporal modification outcomes may need to be refined when applying RMS to other dance styles, such as partner-based dances or styles that place less emphasis on kinematic configuration [50, 51]. Quantifying how RMS maps to participant-reported difficulty and engagement could enable the selection of therapy parameters to optimize the level of challenge during therapy [3, 6, 52]. Future work should relate RMS performance to therapy efficacy, participant-reported difficulty and engagement, and therapy impacts of motor and/or cognitive function [6]. Ultimately, to establish RMS as a clinical tool to help clinicians optimize dance-based therapies, we envision a unifying multivariate model capable of predicting RMS performance across participants of different ages, cognitive function levels, prior relationships to music or dance, and rhythmic proficiency.

## Conclusions

We investigated the relationships between rhythmic movement sequence (RMS) performance and novel assessments of individuals’ relationships to music and dance, as well as rhythmic proficiency. The associations of only music relationships and rhythmic proficiency with temporal RMS in young and older adults without, but not with, MCI suggest that cognitive deficits in adults with MCI likely hinder the ability of music relationships or rhythmic proficiency to improve performance on dance-like RMS. These findings contribute to a growing understanding of the factors influencing the ability to accurately perform dance-like movements, which may inform the personalization of dance-based therapies.

## Supporting information

Rhythm Assessment

Assessment analysis table

Data dictionary

## Author contributions

Alexandra Slusarenko (Conceptualization; Formal Analysis; Investigation; Visualization; Writing – original draft; Writing – review & editing)

Michael C. Rosenberg (Conceptualization; Data Curation; Funding Acquisition; Investigation; Methodology; Project administration; Software; Validation; Visualization; Writing – original draft; Writing – review & editing)

Meghan E. Kazanski (Conceptualization; Investigation; Methodology; Project administration; Validation; Writing – review & editing)

J. Lucas McKay (Conceptualization; Data Curation; Funding Acquisition; Methodology; Software; Supervision; Writing – review & editing)

Laura Emmery (Conceptualization; Funding Acquisition; Methodology; Supervision; Writing – review & editing)

Trisha M. Kesar (Conceptualization; Funding Acquisition; Methodology; Supervision; Writing – review & editing)

Madeleine E. Hackney (Conceptualization; Data Curation; Funding Acquisition; Methodology; Project administration; Resources; Supervision; Writing – review & editing)

## Acknowledgments

We thank K. Cao, T. Prusin, and C. Carroll-Sauer for their assistance in data collection and participant recruitment.

## Data availability

The data supporting the findings of this study are openly available as Supplemental Material. These data were derived from the following resources available in the public domain: https://www.frontiersin.org/articles/10.3389/fnhum.2023.1040930/full.

## References

[1] Zhu Y, Zhong Q, Ji J, Ma J, Wu H, Gao Y, Ali N, Wang T (2020) Effects of aerobic dance on cognition in older adults with mild cognitive impairment: a systematic review and meta-analysis. J Alzheimers Dis 74, 679–690.

[2] Gauthier S, Reisberg B, Zaudig M, Petersen RC, Ritchie K, Broich K, Belleville S, Brodaty H, Bennett D, Chertkow H, Cummings JL, de Leon M, Feldman H, Ganguli M, Hampel H, Scheltens P, Tierney MC, Whitehouse P, Winblad B (2006) Mild cognitive impairment. The Lancet 367, 1262–1270.

[3] Lazarou I, Parastatidis T, Tsolaki A, Gkioka M, Karakostas A, Douka S, Tsolaki M (2017) International ballroom dancing against neurodegeneration: a randomized controlled trial in Greek community-dwelling elders with mild cognitive impairment. Am J Alzheimers Dis Other Demen 32, 489–499.

[4] Zhu Y, Wu H, Qi M, Wang S, Zhang Q, Zhou L, Wang S, Wang W, Wu T, Xiao M (2018) Effects of a specially designed aerobic dance routine on mild cognitive impairment. Clin Interv Aging 13, 1691.

[5] McKee KE, Hackney ME (2013) The effects of adapted tango on spatial cognition and disease severity in Parkinson’s disease. J Mot Behav 45, 519–529.

[6] Guadagnoli MA, Lee TD (2004) Challenge point: a framework for conceptualizing the effects of various practice conditions in motor learning. J Motor Behav 36, 212–224.

[7] Hackney ME, Earhart GM (2009) Effects of dance on movement control in Parkinson’s disease: a comparison of Argentine tango and American ballroom. J Rehabil Med 41, 475–481.

[8] Rosenberg MC, Slusarenko A, Cao K, Lucas McKay J, Emmery L, Kesar TM, Hackney ME (2023) Motor and cognitive deficits limit the ability to flexibly modulate spatiotemporal gait features in older adults with mild cognitive impairment. Front Hum Neurosci 17.

[9] Rallis I, Doulamis N, Doulamis A, Voulodimos A, Vescoukis V (2018) Spatio-temporal summarization of dance choreographies. Computers & Graphics 73, 88–101.

[10] Hackney ME, Kantorovich S, Levin R, Earhart GM (2007) Effects of tango on functional mobility in Parkinson’s disease: a preliminary study. J Neurol Phys Ther 31, 173–179.

[11] Wilson M, Kwon Y-H (2008) The role of biomechanics in understanding dance movement: a review. J Dance Med Sci 12, 109–116.

[12] Thaut MH, Abiru M (2010) Rhythmic auditory stimulation in rehabilitation of movement disorders: a review of current research. Music Percept 27, 263–269.

[13] Thaut M, Kenyon G, Schauer M, McIntosh G (1999) The connection between rhythmicity and brain function. IEEE Eng Med Biol Mag 18, 101–108.

[14] Emmery L, Hackney ME, Kesar T, McKay JL, Rosenberg MC (2023) An integrated review of music cognition and rhythmic stimuli in sensorimotor neurocognition and neurorehabilitation. Ann N Y Acad Sci 1530, 74–86.

[15] Zatorre RJ, Chen JL, Penhune VB (2007) When the brain plays music: auditory–motor interactions in music perception and production. Nat Rev Neurosci 8, 547–558.

[16] Akombo D (2016) The unity of music and dance in world cultures, McFarland & Company, Inc., Jefferson, North Carolina.

[17] Hackney ME, Earhart GM (2010) Recommendations for implementing tango classes for persons with Parkinson disease. Am J Dance Ther 32, 41–52.

[18] Chen JL, Penhune VB, Zatorre RJ (2008) Moving on time: brain network for auditory-motor synchronization is modulated by rhythm complexity and musical training. J Cogn Neurosci 20, 226–239.

[19] Lerdahl F, Jackendoff R (1983) An overview of hierarchical structure in music. Music Percept, 229–252.

[20] Chen JL, Penhune VB, Zatorre RJ (2008) Listening to musical rhythms recruits motor regions of the brain. Cereb Cortex 18, 2844–2854.

[21] London J (2012) Hearing in time: Psychological aspects of musical meter, Oxford University Press.

[22] Mueller SG, Weiner MW, Thal LJ, Petersen RC, Jack C, Jagust W, Trojanowski JQ, Toga AW, Beckett L (2005) The Alzheimer’s disease neuroimaging initiative. Neuroimag Clin 15, 869–877.

[23] Bowie CR, Harvey PD (2006) Administration and interpretation of the Trail Making Test. Nat Protoc 1, 2277–2281.

[24] Vandierendonck A, Kemps E, Fastame MC, Szmalec A (2004) Working memory components of the Corsi blocks task. Br J Psychol 95, 57–79.

[25] Hackney ME, Hall CD, Echt KV, Wolf SL (2013) Dancing for balance: feasibility and efficacy in oldest-old adults with visual impairment. Nurs Res 62, 138–143.

[26] Nasreddine ZS, Phillips NA, Bédirian V, Charbonneau S, Whitehead V, Collin I, Cummings JL, Chertkow H (2005) The Montreal Cognitive Assessment, MoCA: a brief screening tool for mild cognitive impairment. J Am Geriatr Soc 53, 695–699.

[27] Bläsing B, Schack T (2012) Mental representation of spatial movement parameters in dance. Spat Cogn Comput 12, 111–132.

[28] Ivanenko YP, Cappellini G, Dominici N, Poppele RE, Lacquaniti F (2005) Coordination of locomotion with voluntary movements in humans. J Neurosci 25, 7238–7253.

[29] [29] Styns F, van Noorden L, Moelants D, Leman M (2007) Walking on music. Hum Movment Sci 26, 769–785.

[30] Will U, Berg E (2007) Brain wave synchronization and entrainment to periodic acoustic stimuli. Neurosci Lett 424, 55–60.

[31] [31] Van Noorden L, Moelants D (1999) Resonance in the perception of musical pulse. J New Music Res 28, 43–66.

[32] Drake C, Jones MR, Baruch C (2000) The development of rhythmic attending in auditory sequences: attunement, referent period, focal attending. Cognition 77, 251–288.

[33] Lakatos P, Karmos G, Mehta AD, Ulbert I, Schroeder CE (2008) Entrainment of neuronal oscillations as a mechanism of attentional selection. Science 320, 110–113.

[34] Mancini M, Horak FB (2016) Potential of APDM mobility lab for the monitoring of the progression of Parkinson’s disease. Expert Rev Med Devices 13, 455–462.

[35] Washabaugh EP, Kalyanaraman T, Adamczyk PG, Claflin ES, Krishnan C (2017) Validity and repeatability of inertial measurement units for measuring gait parameters. Gait Posture 55, 87–93.

[36] Slusarenko A (2022) in Neuroscience and Behavioral Biology Emory University.

[37] Bengtsson SL, Ullen F, Ehrsson HH, Hashimoto T, Kito T, Naito E, Forssberg H, Sadato N (2009) Listening to rhythms activates motor and premotor cortices. Cortex 45, 62–71.

[38] Hackney ME, Nocera J, Creel T, Riebesell MD, Kesar T (2017) Exercise and balance in older adults with movement disorders In Locomotion and Posture in Older Adults Springer, pp. 323–346.

[39] Hrysomallis C (2011) Balance ability and athletic performance. Sports Med 41, 221–232.

[40] Busquets A, Ferrer-Uris B, Angulo-Barroso R, Federolf P (2021) Gymnastics experience enhances the development of bipedal-stance multi-segmental coordination and control during proprioceptive reweighting. Front Psychol 12, 661312.

[41] Reimann H, Ramadan R, Fettrow T, Hafer JF, Geyer H, Jeka JJ (2020) Interactions between different age-related factors affecting balance control in walking. Front Spor Act Living 2, 94.

[42] Mancini M, King L, Salarian A, Holmstrom L, McNames J, Horak FB (2011) Mobility lab to assess balance and gait with synchronized body-worn sensors. J Bioeng Biomed Sci, 007.

[43] Spain R, George RS, Salarian A, Mancini M, Wagner J, Horak F, Bourdette D (2012) Body-worn motion sensors detect balance and gait deficits in people with multiple sclerosis who have normal walking speed. Gait Posture 35, 573–578.

[44] Morrow MM, Lowndes B, Fortune E, Kaufman KR, Hallbeck MS (2017) Validation of inertial measurement units for upper body kinematics. J Appl Biomech 33, 227–232.

[45] Tulipani L, Boocock MG, Lomond KV, El-Gohary M, Reid DA, Henry SM (2018) Validation of an inertial sensor system for physical therapists to quantify movement coordination during functional tasks. J Appl Biomech 34, 23–30.

[46] Müllensiefen D, Gingras B, Musil J, Stewart L (2014) The musicality of non-musicians: An index for assessing musical sophistication in the general population. PLoS One 9, e89642.

[47] Rose D, Müllensiefen D, Lovatt P, Orgs G (2022) The Goldsmiths Dance Sophistication Index (Gold-DSI): A psychometric tool to assess individual differences in dance experience. Psychol Aesthet Crea 16, 733–745.

[48] Mas-Herrero E, Marco-Pallares J, Lorenzo-Seva U, Zatorre RJ, Rodriguez-Fornells A (2012) Individual differences in music reward experiences. Music Percept 31, 118–138.

[49] Karpati FJ, Giacosa C, Foster NE, Penhune VB, Hyde KL (2015) Dance and the brain: a review. Ann N Y Acad Sci 1337, 140–146.

[50] Hackney ME, Earhart GM (2010) Effects of dance on gait and balance in Parkinson’s disease: a comparison of partnered and nonpartnered dance movement. Neurorehabil Neural Repair 24, 384–392.

[51] Hackney ME, Kantorovich S, Earhart GM (2007) A study on the effects of Argentine tango as a form of partnered dance for those with Parkinson disease and the healthy elderly. Am J Dance Ther 29, 109–127.

[52] Chan JS, Wu J, Deng K, Yan JH (2020) The effectiveness of dance interventions on cognition in patients with mild cognitive impairment: a meta-analysis of randomized controlled trials. Neurosci Biobehav Rev 118, 80–88.

