## Supplemental - S1 for "Associations between music and dance relationships, rhythmic proficiency, and spatiotemporal movement modulation ability in adults with and without mild cognitive impairment"

### README

This file describes the information contained in supplemental files

**Supplemental Media**

**Supplemental Tables**
1. DataDictionary.csv
2. Table_AssessmentItemIndependence.csv
3. Table_Data.csv

**Rhythm Assessment Instructions (S2)**

**Supplemental Results (S3)**
1. Participant preference and engagement in music and dance
2. Correlations between the RA composite scores without the *visual rhythm reproduction* item and RMS performance
3. Correlations between the RA subcomponent composite scores and the overall RA composite score
4. Correlations between the RA subcomponent composite scores and the MRQ and DRQ composite scores

#### Supplemental Media

The supplemental videos contain examples of spatial and temporal modifications.

*SupplementalV1_Tutorial_Spatial_SW_AttitudeDeveloppe.mp4* shows the training video for the *Attitude-Developpe* spatial RMS. This video was used to help participants understand the spatial targets of the modification.

*SupplementalV2_Tutorial_Temporal_SimpleDuple_qqSS.mp4* shows a training video that participants watched to learn how to perform temporal modifications. The video consists of following the instructor in performing the step sequence in the following order: clapping, tapping one foot, shifting weight between the feet, stepping in place, and walking. The sample video is for the *simple duple – qqSS* modification.

*RhythmAssessment.pptx* contains the audio clips and musical pieces used during the Rhythm Assessment. Participants would see these slides as the assessment was being conducted.

### Supplemental Tables

#### DataDictionary.csv

For each supplemental table (the *Domain* column in DataDictionary.csv), DataDictionary.csv contains the following information in each column:

1. Descriptions of general classes of data (*SubDomains*)
2. Each variable's name (*VariableName*)
3. The full, unabbreviated variable name (*FullName*)
4. A description of each variable, including units (*Description*)

#### Table_AssessmentItemIndependence.csv

This table contains Pearson's R coefficients for correlations between each pair of items of the Music Relationship Questionnaire (MRQ), Dance Relationship Questionnaire (DRQ). Correlations were performed between all items within each assessment. Correlations were used to determine the extent to which questionnaire and assessment items covaried. Because each questionnaire and assessment had 10 items, each row contains a generic list of items 1-10. The columns contain items 1-10 specific to the MRQ, DRQ, and RA. For example, the [row, column] pair [*Item_1*, *MRQ_2*] contains the correlation between Items 1 and 2 on the MRQ. *Similarly,* pair [*Item_1*, *RA_7*] contains the correlation between Items 1 and 7 on the RA.

#### Table_Data.csv

This table contains demographic information, clinical assessment scores, composite scores for the MRQ, DRQ, and RA, and percent errors for spatial, temporal, and spatiotemporal rhythmic movement sequences (RMS). Rows denote participants, with the first column containing participant IDs. The YA, OA, and MCI abbreviations denote the young adult, older adult without MCI, and older adult with MCI groups. Column descriptions can be found in the DataDictionary.csv table.

### Rhythm Assessment Instructions (S2)

Supplemental – S2 contains the instructions, script, and scoring for the Rhythm Assessment.

### Supplementary Results (S3)

Supplemental – S3 contains additional results that are relevant to the main manuscript but were not a central focus of the study.

1. Participant preference and engagement in music and dance
2. Correlations between the RA composite scores without the *visual rhythm reproduction item* and RMS performance
3. Correlations between the RA subcomponent composite scores and the overall RA composite score
4. Correlations between the RA subcomponent composite scores and the MRQ and DRQ composite scores

#### Participant preference and engagement in music and dance

To assess participants' preferences for different music genres and the extent of their active engagement in music and dance, we included multiple-choice and open-ended questions, respectively, in the Music Relationship Questionnaire (MRQ) and Dance Relationship Questionnaire (DRQ). This analysis characterizes genres and evaluates the extent to which genre preference impacts RMS performance.

#### Correlations between the RA composite scores without the *visual rhythm reproduction* item and RMS performance

To determine if study conclusions about the relationships between RA composite scores and participants' abilities to read musical notation were impacted by some participants' inability to read musical notation, we repeated RA regression analyses with the visual rhythm reproduction item removed from the RA composite score.

#### Correlations between the RA subcomponent composite scores and the overall RA composite score

To characterize the auditory subcomponents of the RA, we regressed the RA composite scores against auditory rhythm reproduction composite scores and auditory meter recognition scores separately.

#### Correlations between the RA subcomponent composite scores and the MRQ and DRQ composite scores

To characterize the relationship between auditory subcomponents of the RA and composite scores on the MRQ and DRQ, we regressed the MRQ and DRQ composite scores against auditory rhythm reproduction composite scores and auditory meter recognition scores separately.
