## Supplemental - S2 for "Associations between music and dance relationships, rhythmic proficiency, and spatiotemporal movement modulation ability in adults with and without mild cognitive impairment"

### Rhythm Assessment Instructions

The following contains instructions for administering the Rhythm Assessment. A scoring sheet is included in Table S1.

The rhythms and instructions are also included in the *RhythmAssessment.pptx* PowerPoint document.

**Notes for administering the Rhythm Assessment virtually (e.g., via Zoom)**:

1. If administering the assessment virtually (e.g., via Zoom), check your audio settings: Set “Suppress Background Noise” to “Low”.
2. If administering the assessment virtually, record Parts 1 & 2 of the assessment. After the assessment, record a voice memo of the participant’s clapping in Parts 1 & 2.
3. Test a random rhythm to ensure that the participant can hear the audio.

In the following sections, instructions spoken to the participant are *italicized*.

#### Introduction

“*There are three parts to this assessment. During the first part, you will be listening and clapping. During the second part, you will be reading musical notation and clapping. During the third part, you will just be listening. I will record your clapping as a voice memo*.”

#### Part 1: Auditory rhythm reproduction

Each of four rhythms is played three times. After the third time that a rhythm is played, the participant will attempt to clapback the rhythm twice. The order of rhythms is not randomized.

“*Just to be clear, you will hear each rhythm one at a time, and I will play that rhythm three times. Then you will clap back twice. Are you ready*?”

You can now begin playing the first rhythm.

**Note**: Indicate the number of ”playings” after each rhythm is played. For example, “First playing: *rhythm 1*, Second playing: *rhythm 1*, and Third playing: *rhythm 1*”

After the rhythm plays for the third time, ask the participant to clapback the rhythm twice. Then proceed to the next rhythm.

**Scoring**:
Correct = 1
Partially correct (one of two clapbacks is correct) = 0.5
Incorrect = 0

#### Part 2: Visual rhythm reproduction

The participant will attempt to read and clapback musical notes.
“*You may practice and let me know when you are ready.*”

**Note**: If a participant does not have a musical background, but wants to attempt this part anyway, make a note of this and let the participant proceed.

Ask and note if each participant has a musical background AFTER they have clapped and read the rhythm.

**Scoring**:
Correct: 1
Incorrect: 0

#### Part 3: Meter recognition

The participant will listen to music clips and attempt to identify if the rhythm is in "twos" (Duple), "threes" (Waltz), or "other." The correct answers are listed in the **Table S1** scoring sheet, below.

"*Just to be clear, you will hear each clip one at a time. Each clip is about 30s. You have up to three times to hear it played, and you may give your final answer after the first or second playing. Your answer choices are twos, threes, or other. Are you ready?*"

**Part 3 Key**:

a. Twos
b. Threes
c. Twos
d. Threes
e. Other

**Scoring**:
Correct: 1
Incorrect: 0

**Table S1**: RA scoring sheet.

| Item | Item Description | Scoring Scale – Response (Points) | | |
|---|---|---|---|---|
| | | Correct | Partially correct | Incorrect |
| 1 | Auditory rhythm reproduction A | | | |
| 2 | Auditory rhythm reproduction B | | | |
| 3 | Auditory rhythm reproduction C | | | |
| 4 | Auditory rhythm reproduction D | | | |
| 5 | Visual rhythm reproduction | | - | |
| 6 | Auditory meter recognition A (Duple) | (twos) | - | |
| 7 | Auditory meter recognition B (Waltz) | (threes) | - | |
| 8 | Auditory meter recognition C (Duple) | (twos) | - | |
| 9 | Auditory meter recognition D (Waltz) | (threes) | - | |
| 10 | Auditory meter recognition E (Other) | (other) | - | |

"Partially correct" indicated that the participant clapped the rhythm correctly once and incorrectly once.
