## Supplemental - S3 for "Associations between music and dance relationships, rhythmic proficiency, and spatiotemporal movement modulation ability in adults with and without mild cognitive impairment"

Overview: This supplemental file describes the information contained in supplemental data tables.

### Participant preference and engagement in music and dance

To assess participants' preferences for different music genres and the extent of their active engagement in music and dance, we included multiple-choice and open-ended questions, respectively, in the Music Relationship Questionnaire (MRQ) and Dance Relationship Questionnaire (DRQ). Supplemental Table 1 shows the MRQ and DRQ music and dance styles for multiple-choice questions.

**Supplemental Table 1**: Multiple-choice questions and choices in the MRQ and DRQ

| **MRQ: "What kind(s) of music do you listen to?"** | **DRQ: "What kind(s) of dance do you mostly do, if any?"** |
|---|---|
| Classical | Ballet |
| Rock | Ballroom |
| Pop | Contemporary |
| Hip hop | Hip hop |
| Rap | Jazz |
| Christian Gospel | Tap |
| R&B | Folk |
| Jazz | Irish |
| Heavy Metal | Modern |
| Country | Swing |
| Folk | Other |
| Blues | |
| Electronic | |
| Other | |

**MRQ open-ended question**: "Do you create your own music (compose, improvise, DJ, etc.)? If so, tell us what you do?"

**DRQ open-ended question**: "Do you create your own dances (choreograph, improvise, teach, perform, etc.)? If so, tell us what you do?"

Below, we provide a summary of participant responses to these questions.

First, to determine if young adults (YA), older adults without MCI (OA), and older adults with MCI (MCI) different in the breadth of preferred musical genres and dance styles, we computed the number of genres and styles that each participant reported listening to or dancing. We tested for differences in breadth of music and dance preference between the YA, OA, and MCI groups using Kruskal-Wallis tests ($\alpha = 0.05$). For each group, we also computed the percent of participants who reported listening to each music genre and doing each dance style.

The median number of music genres listened to was 6, 5, and 3 for the YA, OA, and MCI groups, respectively (Supplementary Figure 1A). However, there was substantial variability in the number of genres, ranging from 1 to 10 across all participants. Musical genres did not differ between groups ($p = 0.379$). Participants reported engaging in fewer dance styles: the median number of dance styles performed was 0, 1, and 1 for the YA, OA, and MCI groups, respectively (range = 0-5; Supplementary Figure 1). The number of dance styles that participants performed did not differ between groups ($p = 0.140$).

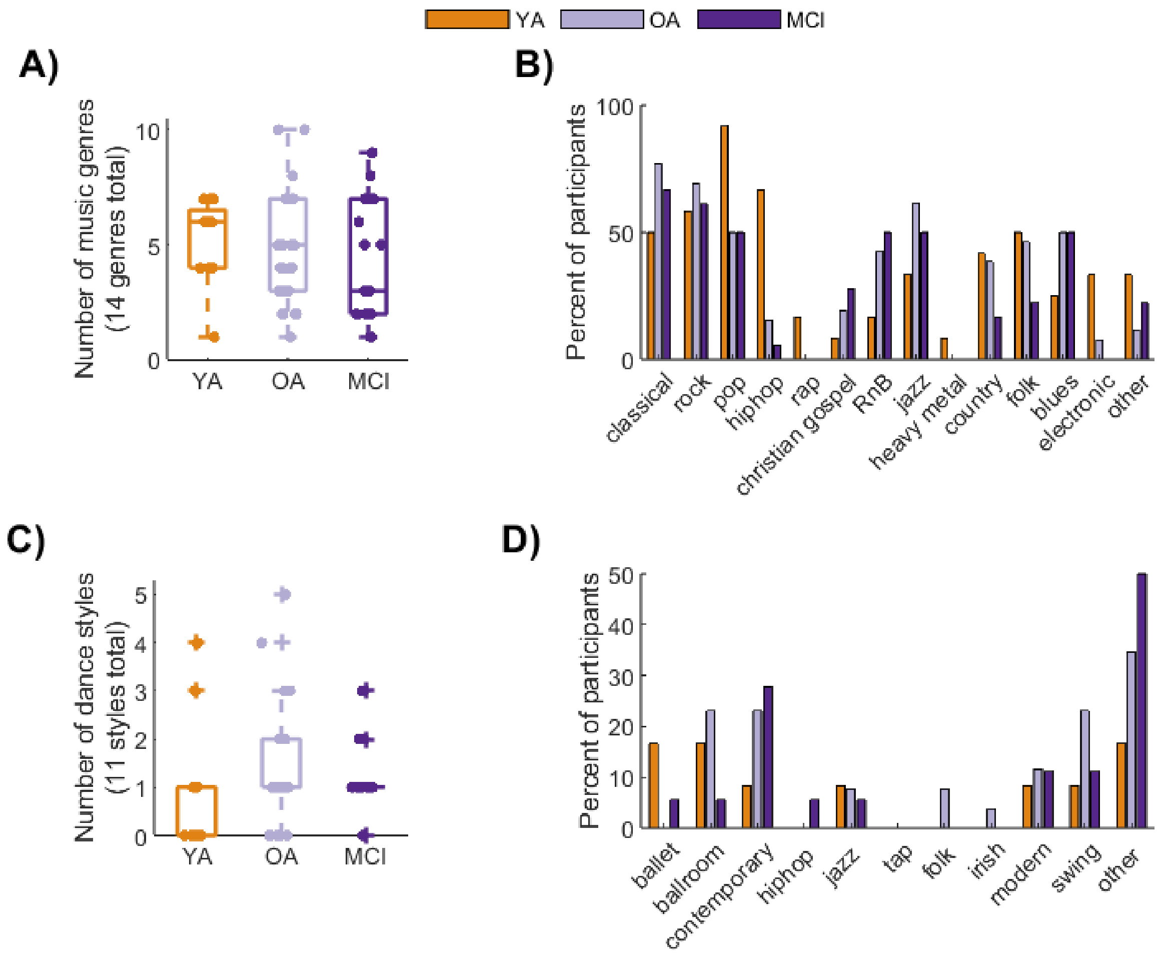


**Supplementary Figure 1: Summary of participant preferences for different music genres and dance styles.** Colors denote the YA (orange), OA (gray) and MCI (purple) groups. A) Percent of music genres, out of 14 total genres, that each participant reported listening to. B) Bar graphs showing the percent of participants that reported listening to each of the 14 music genres. C) Percent of dance styles, out of 11 total styles, that each participant reported performing. D) Bar graphs showing the percent of participants that reported performing each of the 11 dance styles.

Supplementary Figure 1B & D shows the percent of participants that reported listening to each music genre or performing each dance style, respectively. Few notable group differences emerged. However, fewer participants reported engaging in dance than in music. Further, only three participants reported Ballet dancing, which was the basis for RMS spatial modifications.

These results highlight the range of preferences for musical genres and dance styles within our participant cohort. Substantial within-group variability in the breadth of preferences supports future work characterizing how individual-specific music and dance preferences impact RMS performance or therapy efficacy.

Next, we sought to determine if participants who listened to classical music or performed ballet—the music genre and dance style used in this study, respectively—performed better on RMS. If classical music listening or ballet improve RMS performance, then we would expect that participants who engaged in those styles would exhibit better performance on temporal and spatial RMS, respectively. To test this prediction, we computed the percent of participants who both listened to classical music or did ballet and were *top performers* on spatial and temporal RMS. We defined "*top performers*" in each group as the participants with the lowest 50% of RMS percent errors.

Supplementary Figure 2 shows spatial and temporal RMS performance with participants who did ballet (A) and listened to classical music (B) highlighted as squares. Only 2 YA participants and one participant with MCI reported doing ballet. In the YA group, only one of these participants was a *top performer* on spatial RMS (left). The participant with MCI was a *top* performer. All three participants were top *performers* on temporal RMS (right). Conversely, 6 (50%), 20 (77%), and 12 (67%) participants in the YA, OA, and MCI groups reported listening to classical music (also see Supplementary Figure 1B). For spatial RMS in the YA, OA, and MCI groups, 50, 100, 66% of *top-performing* participants reported listening to classical music. For temporal RMS in the YA, OA, and MCI groups, 50, 92, 56% of *top-performing* participants reported listening to classical music.

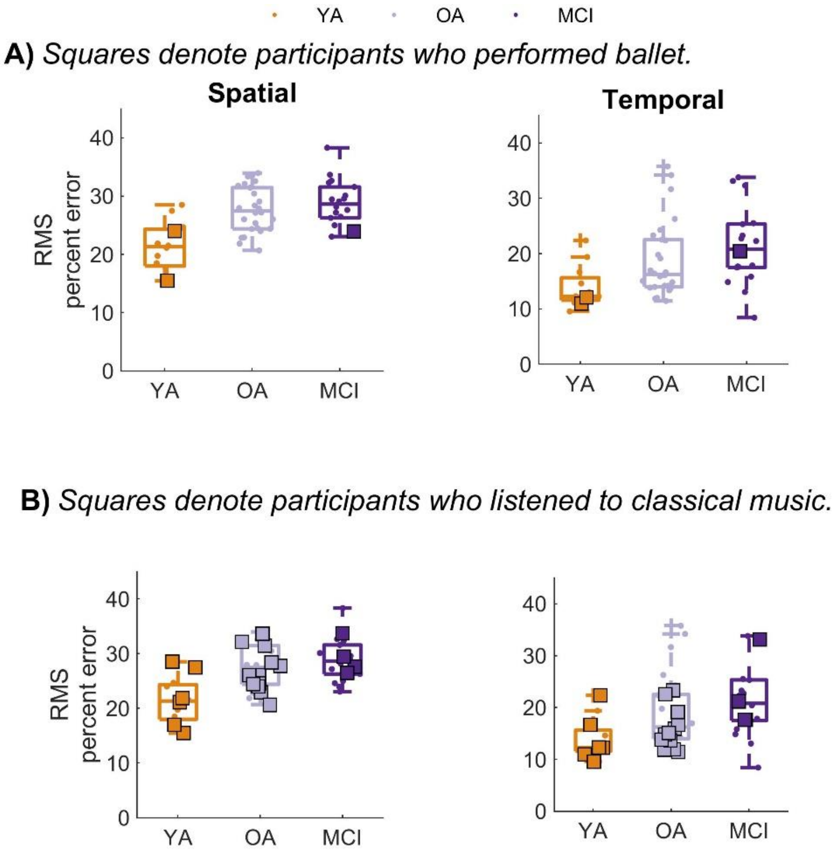


**Supplementary Figure 2: Summary of the relationship ballet performance (A) or classical music listening (B) and RMS performance.** Columns denote the different RMS classes. Colors denote the YA (orange), OA (gray) and MCI (purple) groups. Squares denote participants who reported that they perform ballet (A) or listen to classical music (B). Dots denote participants who reported that they did not perform ballet (A) or listen to classical music (B).

Next, we performed a similar analysis to determine if people who expressed active engagement in music or dance performed better on spatial and temporal RMS. We used an identical median-split approach to determine if the top performers were also those who actively engaged in music or dance, according to responses to the open-ended questions. Any response indicating a level of active engagement was considered to actively engage in music or dance, regardless of genre, style, or level of engagement.

In the YA, OA, and MCI groups only 3, 5, and 1 participants, respectively, reported creating their own music. Similarly, only 4, 6, and 2 participants, respectively, reported creating their own dances. However, YA and OA participants who reported creating their own music or dances (Supplemental Figure 3; squares in the middle column); appeared to perform better (lower error) on temporal RMS than a subset of participants who did not report creating music or dances (Supplemental Figure 3; dots). This was not the case for spatial or spatiotemporal modifications, in which participants who reported creating music or dances were among the worst performers (Supplemental Figure 3; left and right columns).

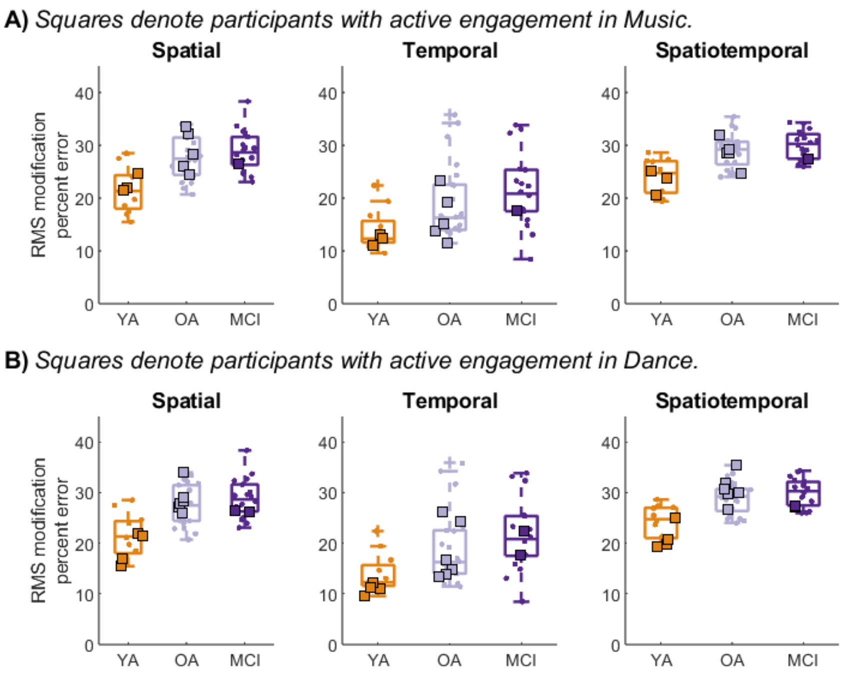


**Supplemental Figure 3: Summary of the relationship between participant engagement in music (A) or dance (B), in relation to RMS performance**. Columns denote the different RMS classes. Colors denote the YA (orange), OA (gray) and MCI (purple) groups. Squares denote participants who reported that they create their own music (A) or dances (B). Dots denote participants who reported that they did not create their own music (A) or dances (B).

These findings support a relationship between classical music listening and RMS performance. Due to the small number of participants who did ballet, we could not determine if ballet performance is likely related to better RMS performance. However, classical music listening appears to be related to better RMS performance. For comparison, if participants were randomly assigned to be *top performers*, we would expect 25, 39, and 33% of participants to be top performers. Therefore, the percent of participants who listen to classical music and were *top performers* on RMS is greater than chance. This relationship is not necessarily unique to classical music; we could provide the same conclusion using (e.g.,) rock, jazz, or folk. Rather than classical music listening impacting RMS performance, performance may be similarly influenced by engagement in a wide array of musical types.

### Correlations between the RA composite scores without the *visual rhythm reproduction* item and RMS performance

To determine if study conclusions about the relationships between RA composite scores and participants' abilities to read musical notation were impacted by some participants' inability to read musical notation, we repeated RA regression analyses with the *visual rhythm reproduction* item removed from the RA composite score. For all participants, we recomputed RA composite scores with the visual rhythm reproduction (*i.e.*, note reading) item removed. As done in the main manuscript, we compared RA scores between the YA and OA groups and between the OA and MCI groups using independent-samples t-tests (α = 0.05). We then regressed the new RA composite scores against temporal and spatiotemporal RMS performance for the YA, OA, and MCI groups separately.

Removing the *visual rhythm reproduction* item from the RA did not substantially change our main results. As in the main manuscript, the RA differed between both the YA and OA groups and the OA and MCI groups (Supplemental Figure 4A). Regression analyses between the RA without the *visual rhythm reproduction* item only slightly altered the regression fits (e.g., for YA temporal RMS, $r^2$ = 0.37 versus 0.40 in the main manuscript; Supplemental Figure 4B & C)

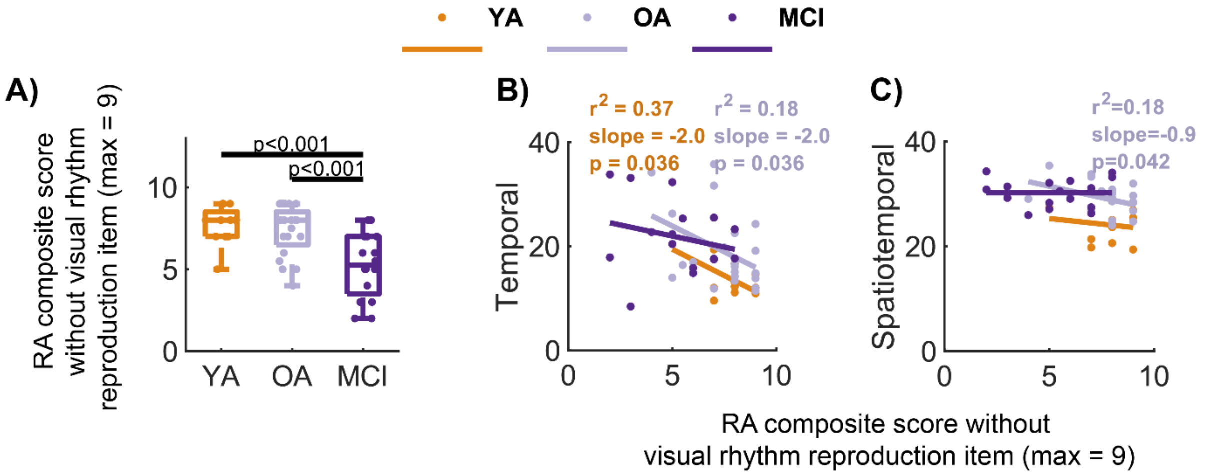


**Supplemental Figure 4: Analysis of RA composite scores excluding the visual rhythm reproduction item.** Colors denote the YA (orange), OA (gray), and MCI (purple) groups. Group-specific regression results report coefficient of determination ($r^2$), slope, and p-value testing that the slope differs from zero (Wald tests; α = 0.05). Regression results are shown only when the regression slopes were significantly different from zero. RMS is compared to A) RA scores compared between groups, B) RA scores regressed against temporal RMS. C) RA scores regressed against spatiotemporal RMS.

These results suggest that the inclusion of the *visual rhythm reproduction* item in the RA did not alter our conclusions.

### Correlations between the RA subcomponent composite scores and the overall RA composite score

To characterize the auditory subcomponents of the RA, we regressed the RA composite scores, against *auditory rhythm reproduction* and *auditory meter recognition* composite scores separately.

For all groups, *auditory rhythm reproduction* composite scores were associated with the overall RA composite scores ($r^2 > 0.38$; $p < 0.001$; Supplemental Figure 5, left). Similarly, for all groups, *auditory rhythm reproduction* composite scores were associated with the overall RA composite scores ($r^2 > 0.48$; $p \leq 0.010$; Supplemental Figure 5, right).

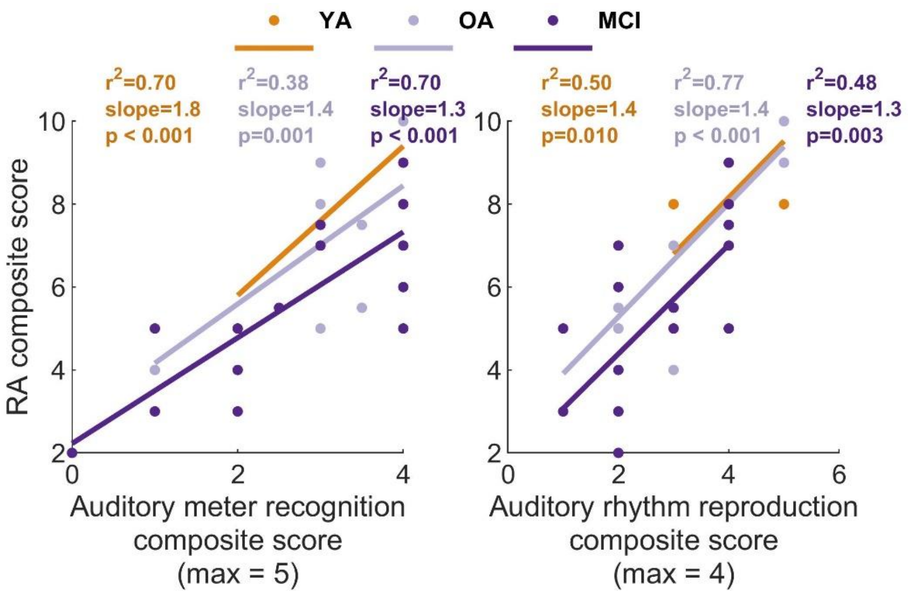


**Supplemental Figure 5: Analysis of RA subcomponent composite scores.** Colors denote the YA (orange), OA (gray), and MCI (purple) groups. Group-specific regression results report coefficient of determination ($r^2$), slope, and p-value testing that the slope differs from zero (Wald tests; $\alpha = 0.05$). Regression results are shown only when the regression slopes were significantly different from zero. Left: RA composite scores regressed against *auditory rhythm reproduction* composite scores. Right: RA composite scores regressed against *auditory meter recognition* composite scores.

The strong relationships between the overall RA composite scores and both the *auditory rhythm reproduction* and *auditory meter recognition* composite scores were not surprising, and suggest that the overall RA scores are not dominated by one subcomponent of the RA.

### Correlations between the RA subcomponent composite scores and the MRQ and DRQ composite scores

To characterize the relationship between auditory subcomponents of the RA and composite scores on the MRQ and DRQ, we regressed the MRQ and DRQ composite scores against auditory rhythm reproduction composite scores and auditory meter recognition scores separately.

For the MRQ, only *auditory rhythm reproduction* composite scores of the RA in the OA group were associated with MRQ composite scores ($r^2 = 0.28$; slope = 0.6; $p = 0.008$; Supplemental Figure 6, top). The *auditory meter recognition* composite scores were not associated with MRQ composite scores in any group. Neither the *auditory rhythm reproduction* nor the *auditory meter recognition* composite scores were associated with DRQ composite scores for any group (Supplemental Figure 6, bottom). However, for all comparisons, the highest scoring participants on the MRQ or DRQ had high *auditory rhythm reproduction* and *auditory meter recognition* composite scores. Similarly, very low scoring participants on the MRQ and DRQ also scored very low on the RA subcomponents.

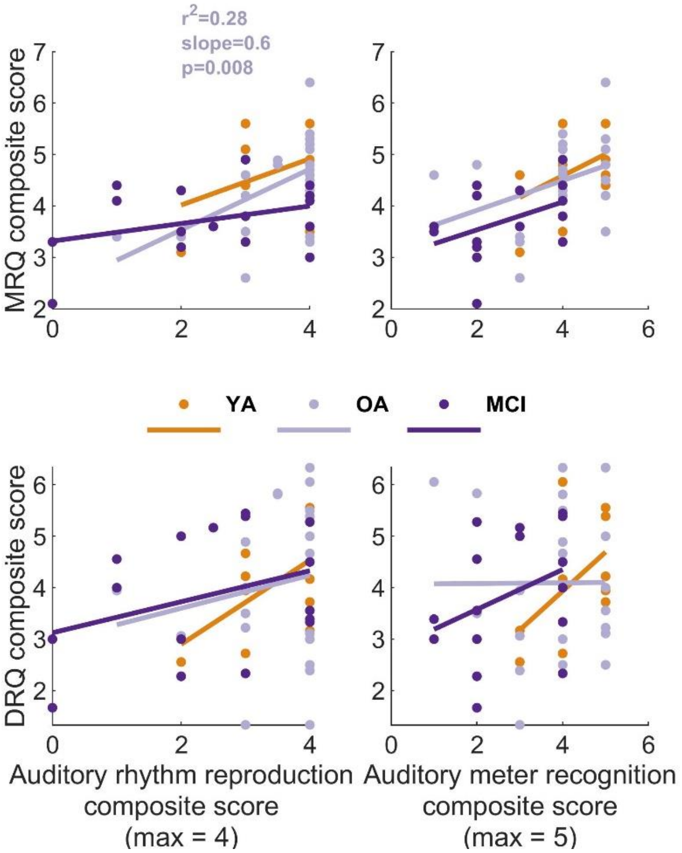


**Supplemental Figure 6: Regression of RA subcomponent composite scores against MRQ (top) and DRQ (bottom) composite scores.** Colors denote the YA (orange), OA (gray), and MCI (purple) groups. Group-specific regression results report coefficient of determination ($r^2$), slope, and p-value testing that the slope differs from zero (Wald tests; $\alpha = 0.05$). Regression results are shown only when the regression slopes were significantly different from zero. Left: MRQ and DRQ composite scores regressed against the *auditory rhythm reproduction* component of the RA. Right: MRQ and DRQ composite scores are regressed against the *auditory meter recognition* component of the RA.

While seldom significant, the highest and lowest scoring participants suggest that both RA subcomponents may be weakly related to MRQ and DRQ scores. The RA subcomponents had only 4 and 5 items, such that we did not expect strong regression results. Higher dimensional assessments may reveal stronger relationships between measures of music or dance relationships and subcomponents of rhythmic proficiency.
