## Supplementary material for "Associations between music and dance relationships, rhythmic proficiency, and spatiotemporal movement modulation ability in adults with and without mild cognitive impairment": Rhythm Assessment

### Slide 1
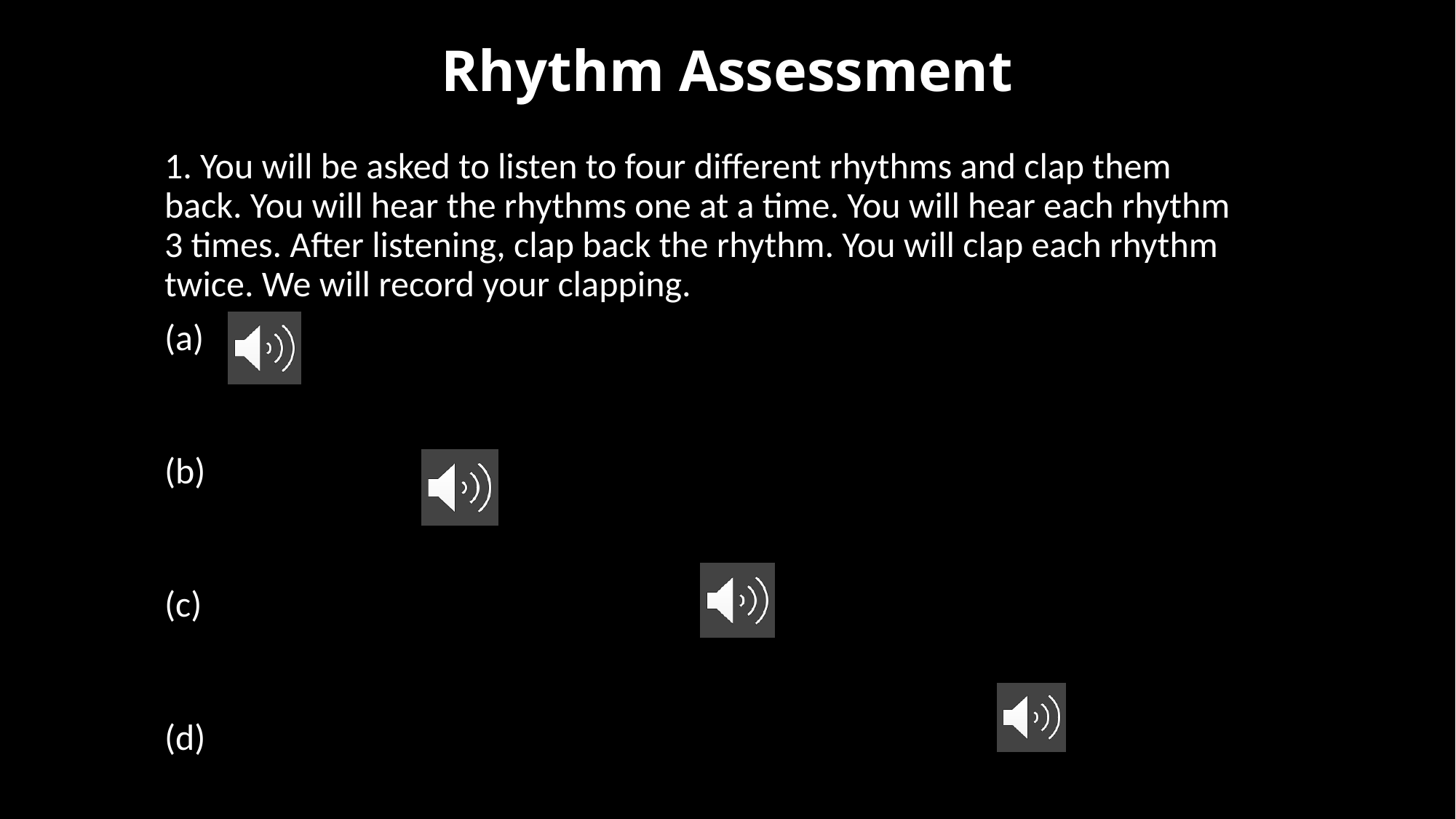

#### Rhythm Assessment
1. You will be asked to listen to four different rhythms and clap them back. You will hear the rhythms one at a time. You will hear each rhythm 3 times. After listening, clap back the rhythm. You will clap each rhythm twice. We will record your clapping.
(a)
(b)
(c)
(d)

### Slide 2
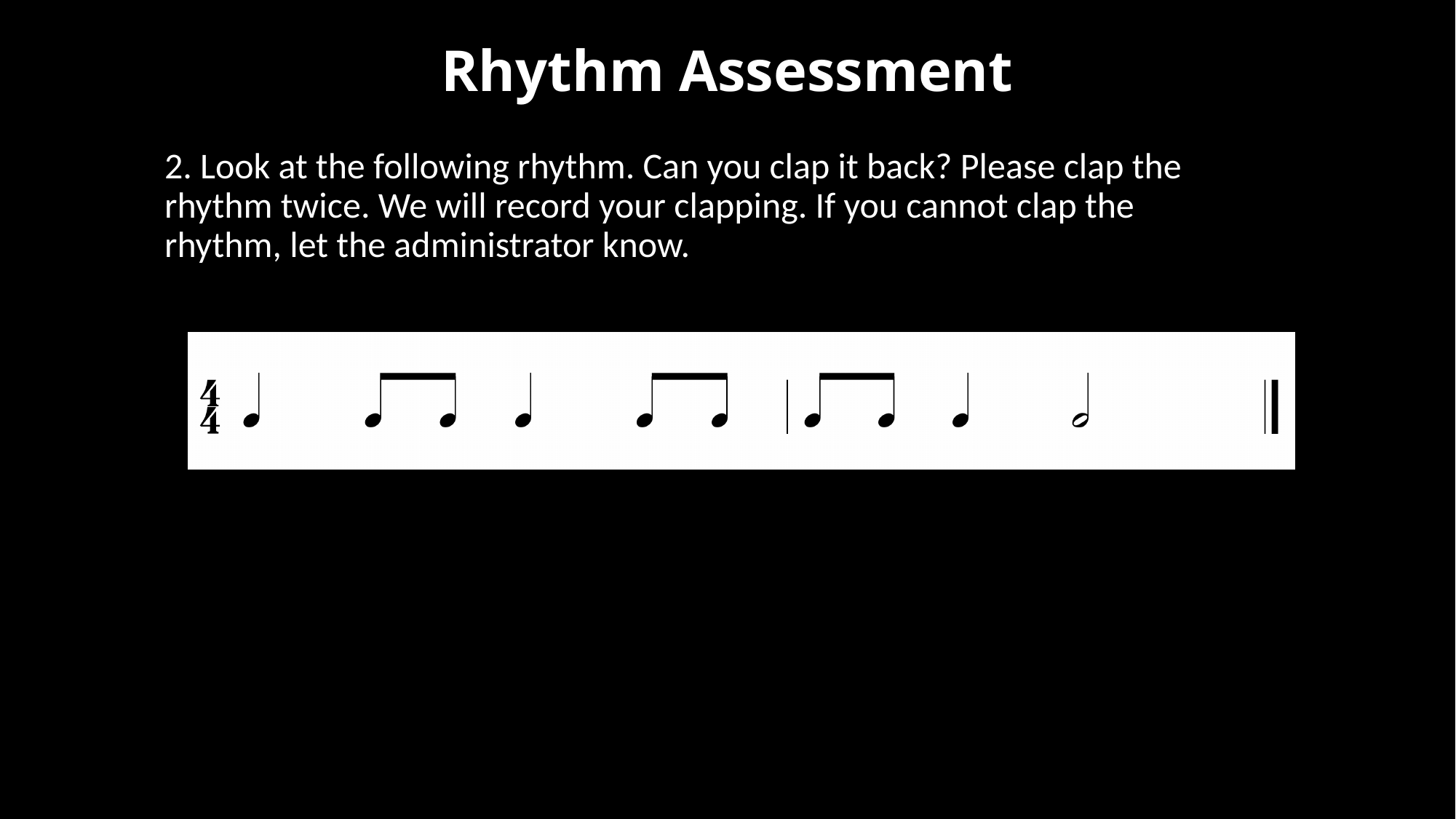

#### Rhythm Assessment
2. Look at the following rhythm. Can you clap it back? Please clap the rhythm twice. We will record your clapping. If you cannot clap the rhythm, let the administrator know.
   ” if they cannot read notation

### Slide 3
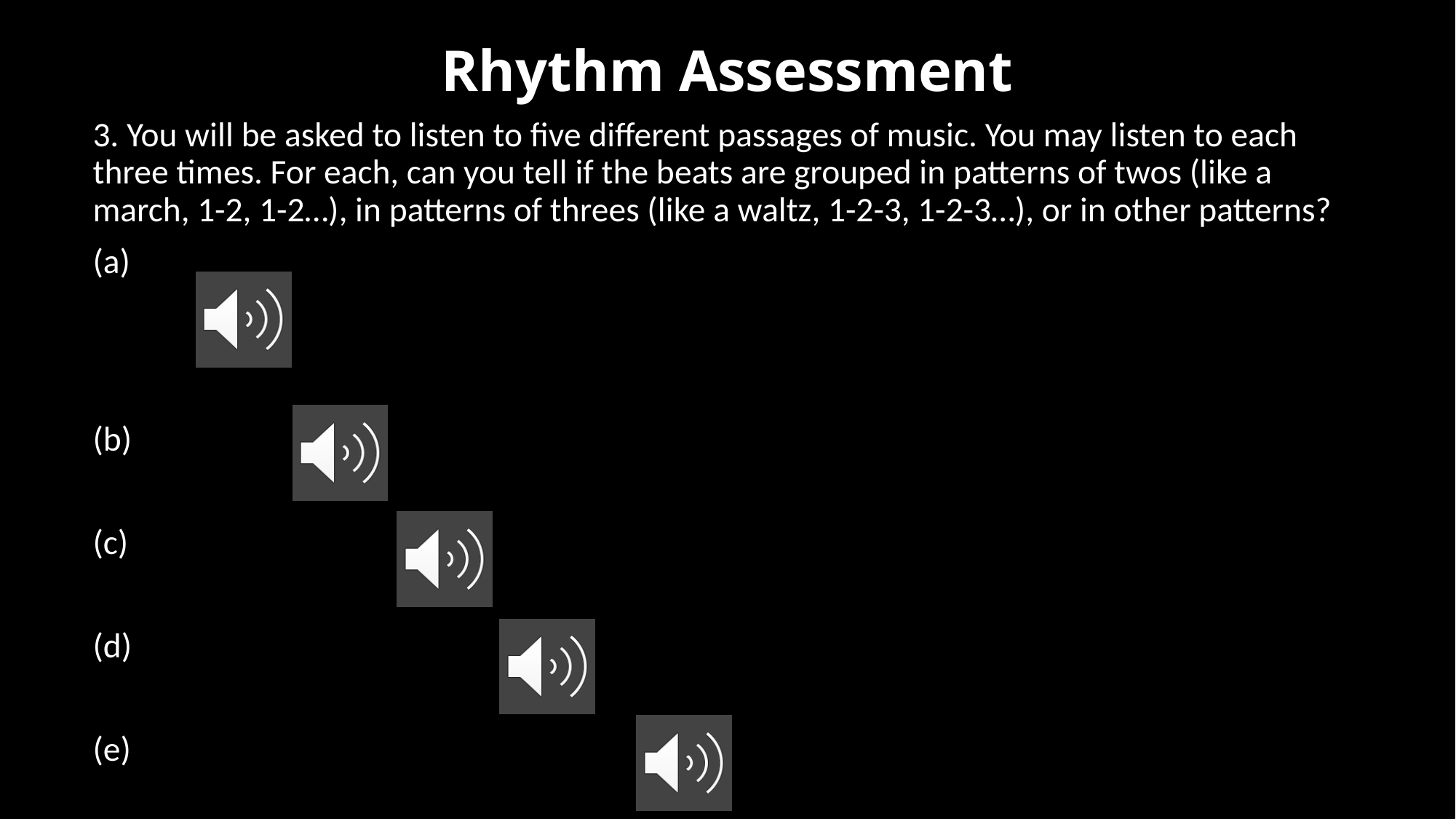

#### Rhythm Assessment
3. You will be asked to listen to five different passages of music. You may listen to each three times. For each, can you tell if the beats are grouped in patterns of twos (like a march, 1-2, 1-2…), in patterns of threes (like a waltz, 1-2-3, 1-2-3…), or in other patterns?
(a)
(b)
(c)
(d)
(e)

### Slide 4
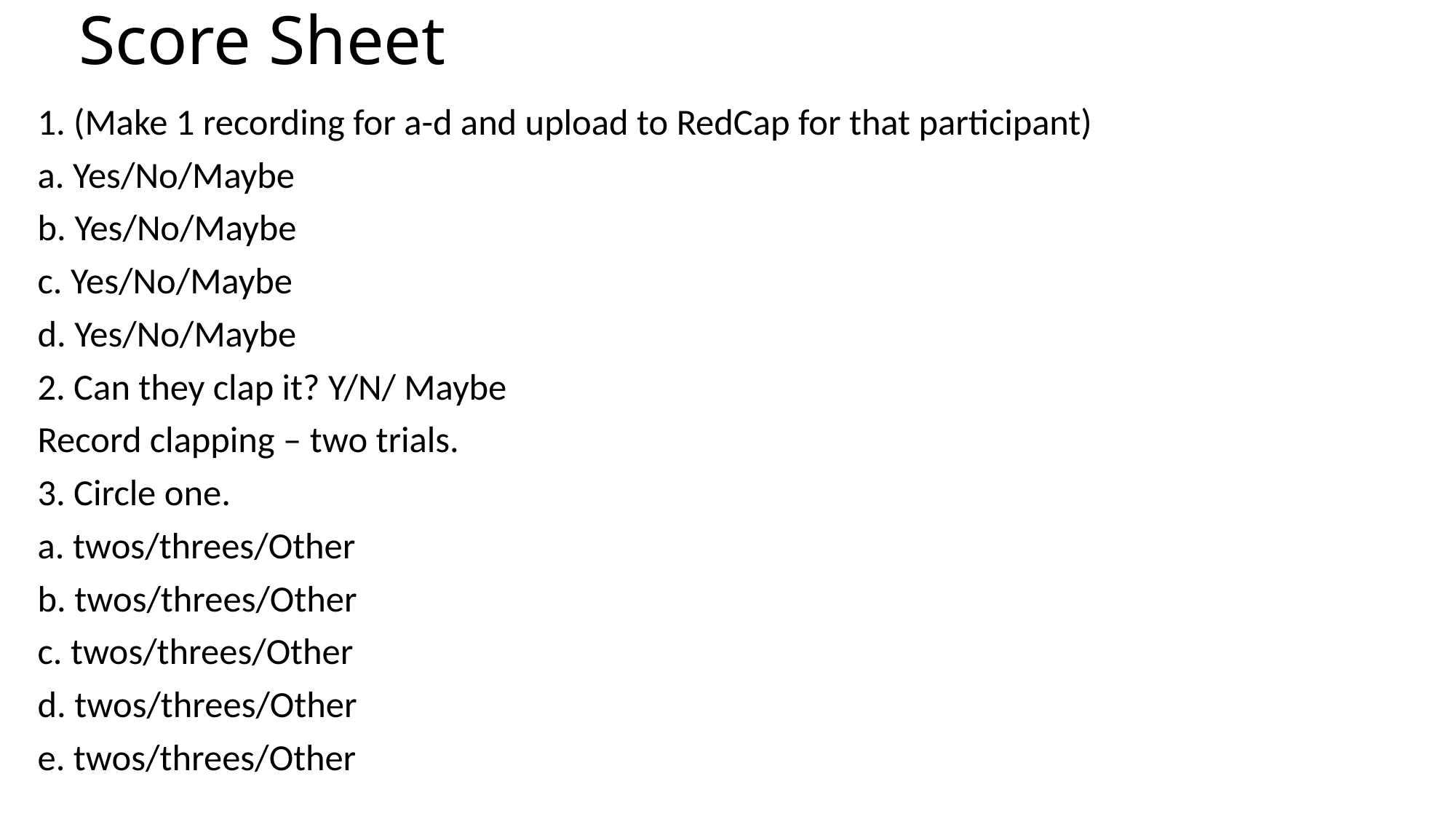

#### Score Sheet
1. (Make 1 recording for a-d and upload to RedCap for that participant)
a. Yes/No/Maybe
b. Yes/No/Maybe
c. Yes/No/Maybe
d. Yes/No/Maybe
2. Can they clap it? Y/N/ Maybe
Record clapping – two trials.
3. Circle one.
a. twos/threes/Other
b. twos/threes/Other
c. twos/threes/Other
d. twos/threes/Other
e. twos/threes/Other
